# Metabolite correlation networks reveal complex phenotypes of adaptation-driving mutations

**DOI:** 10.1101/2025.06.17.660120

**Authors:** Sharon Samuel, Weronika Jasinska, Ezra Sternlicht, Matvey Nikelshparg, Daniel Kleiner, Shai Pilosof, Shimon Bershtein

## Abstract

Living organisms are organized into hierarchical levels of increasing complexity. As mutational effects propagate through these levels, multiple phenotypes are produced. However, identifying which phenotypes influence fitness and drive selection remains a key challenge in understanding genotype-phenotype-fitness relationships. Here, we demonstrate that mutation-induced structural changes in metabolite correlation networks—an organizational level with emergent properties shaped by the cumulative effects of multiple biochemical reactions and regulatory mechanisms—constitute complex phenotypes linking mutations to bacterial fitness. We engineered a metabolically suboptimal *E. coli* strain by replacing the *metK* gene encoding methionine adenosyltransferase (MAT) with an ortholog from *U. urealyticum* and subjected it to laboratory evolution. Analysis of correlation networks constructed from 2,118 untargeted metabolites revealed that adaptive mutations enhanced the strain’s fitness by reducing network density, removing node hubs, and extensively rewiring connectivity. These changes yielded smaller, more cohesive, and better-interconnected network clusters. Moreover, evolution shifted the node representing S-adenosylmethionine (SAM), the product of the MAT-catalyzed reaction, from a peripheral role to a key connector between network clusters with a significantly increased betweenness. None of the accumulated mutations potentially driving this transition directly influenced MAT activity or SAM metabolism, indicating that the shift in SAM’s role is an adaptive phenotype emerging from the metabolic system’s complexity. Targeted metabolomics of key nodes displaying similar network transitions unveiled additional metabolites involved in SAM-related pathways. We propose the construction and analysis of metabolite correlation networks as an experimental and analytical framework for mapping genotype-phenotype-fitness relationships and exploring the mechanisms of metabolic adaptation.

## Introduction

Mapping genotype-phenotype-fitness relationships in evolutionary adaptations is a central problem in evolutionary biology^1^. The primary obstacle lies in the hierarchical organization of biological systems, which spans multiple levels of increasing complexity from individual macromolecules to cell systems, cells, organisms, and populations^2, 3^. A key feature of biological complexity is that each organizational level exhibits unique emergent properties that cannot be fully understood or predicted from the knowledge of its lower-level components^4–6^. While mutations that fuel evolution originate at the genomic level, their phenotypic consequences are shaped by the emergent properties as mutational effects propagate through higher organizational levels^7^. Identifying which among the multiple phenotypes that arise across the levels of complexity influence fitness and ultimately determine the outcome of selection remains a major challenge^1, 8–11^.

Among the cellular systems that shape the phenotypic effects of mutations in bacteria^8^, the metabolic system, charged with the synthesis of building blocks and the production of energy, is most closely linked to organismal fitness. The central role of metabolism in organismal fitness and survival, as evidenced by numerous laboratory evolution studies^10, 12–22^, makes it a critical intermediary in connecting genetic variation with fitness outcomes. Metabolism is carried out by a dense and tightly regulated network of biochemical reactions that catalyze the synthesis of metabolites. When the relative abundance of individual metabolites is measured repeatedly under constant conditions and in the absence of genetic changes, a substantial fraction of detected metabolites exhibits a highly significant correlation between biological replicates^23–27^. The concentration interdependency of metabolites is a unique phenomenon not observed among constituents of other cellular systems, such as mRNA transcripts and proteins, and is rooted in the direct conversion of metabolites into one another via biosynthetic pathways^23^. Two main sources drive metabolite correlations: intrinsic metabolic fluctuations caused primarily by stochastic internal changes in cellular components and pathways^28–32^ and environmental sensitivity, where even micro-perturbations in growth conditions, such as temperature or aeration gradients, affect metabolite levels^25, 33^. Fluctuations in metabolite concentrations tend to propagate beyond neighboring metabolites in a pathway and extend to associated and co-regulated metabolites occupying distant parts of the metabolic system^23, 34^. Thus, metabolite correlation patterns encompass the entire metabolic system and provide insights into its global physiological state under specific genetic and environmental backgrounds.

Correlated metabolites can be represented as networks, where nodes correspond to individual metabolites, and edges represent pairwise correlations^25^. The edges are undirected, indicating mutual correlation without implying causation or directionality, and weighted, reflecting the correlation strength. A key feature of these networks is their dynamic nature; they readily adapt in response to genetic and environmental changes, mirroring the system’s current metabolic state^23, 25^. The metabolite correlation networks constitute, therefore, a complex organizational level characterized by emergent properties originating from the cumulative effects of numerous biochemical reactions and regulatory mechanisms. Consequently, mutation-induced *changes* in network structure and function can be considered complex phenotypic effects. Therefore, we hypothesized that mapping the genetic changes that accumulate in evolving bacterial populations onto the properties of metabolite correlation networks would allow to delineate the complex adaptive phenotypes of mutations and elucidate the genotype-phenotype-fitness relationship driving the evolutionary process.

To explore this hypothesis, we performed adaptive laboratory evolution driven by an inefficient enzyme. In nature, bacterial populations frequently experience novel selection pressures arising from various genetic and ecological factors that produce inefficient enzymatic activities, including random mutations that inactivate specialized enzymes^19, 35–38^, environmental changes that favor secondary, promiscuous enzymatic activities^39, 40^, or horizontal gene transfer events that introduce foreign genes whose enzymatic products are poorly suited to the host’s metabolic needs^22, 41, 42^. Notably, mutations that accumulate in bacterial populations under selection to improve an inefficient enzyme activity only rarely target the gene encoding the inefficient enzyme itself. Instead, they tend to occur in other metabolic and regulatory genes, often causing loss-of-function (LOF) effects^19, 35, 37, 43–45^. The adaptive basis of these off-target LOF mutations is not always clear, especially when they co-occur in multiple genes or involve pleiotropic genes (genes affecting multiple traits), such as those encoding RNA polymerase subunits, translation termination release factors, or metabolic enzymes controlling cellular redox potential^35, 46, 47^.

We reasoned that the fitness effects of mutations accumulating in bacterial populations adapting to an inefficient enzyme can be determined by characterizing structural and functional changes in metabolite correlation network. To model inefficient enzyme evolution in laboratory, we generated and evolved a metabolically suboptimal *Escherichia coli* strain in which the endogenous *metK* gene, encoding an essential methionine adenosyltransferase (MAT), was replaced with an ortholog from *Ureaplasma urealyticum* whose activity is poorly suited to *E. coli*’s metabolic needs^48^. We identified 2,118 untargeted metabolic features in variants isolated from both the evolved and ancestral populations of the modified strain, as well as from the evolved and ancestral wild-type *E. coli*. We then constructed metabolite correlation networks composed exclusively of robust correlations that withstand micro-environmental perturbations. Our analysis showed that network properties at global, meso, and individual node scales link genetic changes acquired during evolution to their phenotypic and fitness effects, even though most adaptive mutations occurred in genes seemingly unrelated to MAT activity or SAM metabolism. Furthermore, by integrating targeted metabolomics, we predicted metabolic identity of selected individual nodes whose roles and properties in the untargeted metabolite correlation networks shifted during adaptive evolution. We found that a substantial fraction of the predicted metabolites participates in SAM-related metabolic pathways and global metabolic regulation, opening new avenues for exploring the mechanisms underlying mutational effects on metabolism.

## Results

### Adaptation to an inefficient enzyme involves off-target pleiotropic mutations that reshape metabolism and enhance fitness

To model inefficient enzyme evolution in laboratory settings, we replaced the chromosomal *metK* gene in *E. coli* with an ortholog from *U. urealyticum,* while retaining the endogenous *E. coli* promoter (**Fig. 1A**). The *metK* gene encodes methionine adenosyltransferase (MAT), an essential enzyme responsible for synthesizing S-adenosylmethionine (SAM) from ATP and methionine^49^. SAM is involved in numerous biochemical processes, including the methylation of DNA, proteins, RNA, and small molecules, the synthesis of enzyme cofactors, and the production of polyamines^50^.

**Figure 1.**
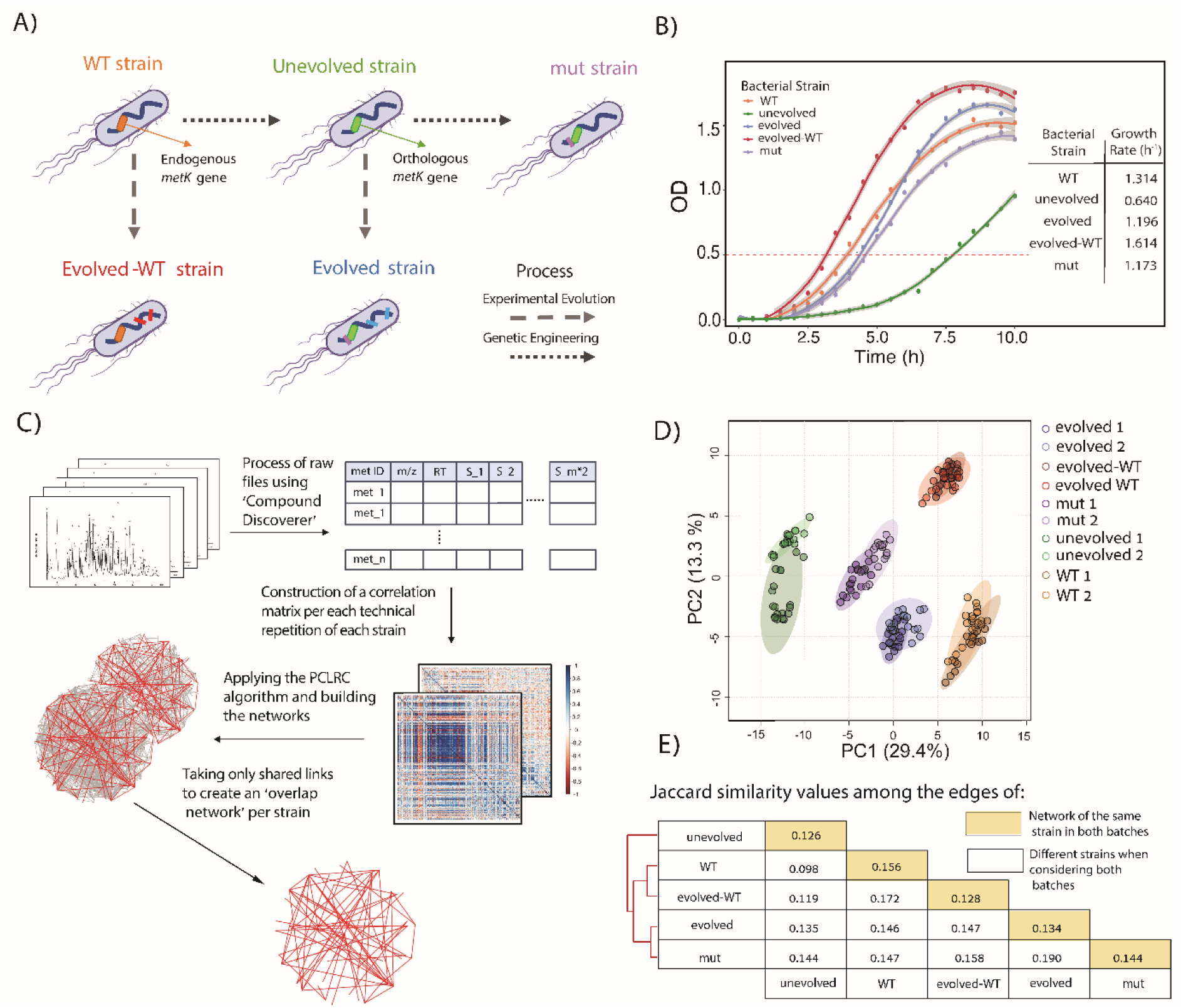
The experimental system. **A.** Gene editing and adaptive laboratory evolution of *E. coli* strains. **B.** Growth curves and growth rates (in 1/hour) of WT, ‘unevolved’, and ‘mut’ strains as well as ‘evolved’ and ‘evolved-WT’ populations. The shaded area represents STD of 4 independent measurements. (**C-E**). Data collection and construction of metabolite correlation networks. **C.** A pipeline for processing untargeted metabolomics data and constructing correlation networks. Raw chromatograms from LC-MS analysis of 19 replicates per batch (across two batches) are processed using Compound Discoverer software (Thermo Scientific), generating data files containing the relative abundances of 2,118 individual metabolic features (Met_1, Met_2, …, Met_n) in the different samples (S_1, S_2,…, S_m*2, when m represents the number of samples in each experimental batch) along with their corresponding mass-to-charge (m/z) ratios and retention times. For each strain, two pairwise correlation matrices (one per batch) are created, representing relative metabolite abundances, and used to construct two correlation networks. These networks are then filtered using the PCLRC algorithm to reduce non-specific (transitive) interactions. Finally, a single network is constructed, containing only the links shared by both batch networks. **D.** Principal component analysis (PCA) of the relative abundances of untargeted metabolic features extracted from both batches. Each data point represents a biological replicate within a batch. Blue arrows indicate the direction of changes in metabolic states induced by genome editing, while red arrows indicate the direction of changes to evolutionary adaptations. **E.** Jaccard similarity analysis and hierarchical clustering of the metabolic networks. Yellow cells indicate similarity between the two batch networks within each strain, while white cells represent similarity between the overlap networks of different strains. The red dendrogram shows unsupervised hierarchical clustering based on the Jaccard similarity values.

While both *E. coli* MAT (EcMAT) and *U. urealyticum* MAT (UuMAT) display similar enzymatic constants, UuMAT is substantially more sensitive to SAM inhibition, and its catalytic efficiency drops by more than half in the presence of the intracellular SAM levels of *E. coli*^48^. The *E. coli* strain with the orthologous *metK* replacement (dubbed the ‘unevolved’ strain) showed a severe growth rate reduction from 1.3 h^-^¹ to 0.64 h^-^¹ (One-way ANOVA across strains: F4,5 =1133, *p-value* =1. 41·10^-7^, followed by Tukey’s HSD, *p-value* < 0.001) (**Fig. 1B, Table S1**), a metabolic imbalance characterized by a significantly decreased SAM and increased methionine levels (One-way ANOVA across strains: SAM, F4,185 = 168.2, *p-value* < 2·10^-16^, followed by Tukey’s HSD, *p-value* < 2·10^-7^; methionine, F4,185 = 434.6, *p-value* <2·10^-16^, followed by Tukey’s HSD, *p-value* < 2·10^-7^) (**Fig. S1A**), and an 80% decrease in the intracellular MAT abundance (**Fig. S1B**).

We subjected both the ‘unevolved’ and wild-type (WT) *E. coli* strains to a laboratory evolution by serial passaging for the duration of 320 generations, resulting in the ‘evolved’ and ‘evolved-WT’ populations, respectively (**Fig. 1A).** The growth rate of the ‘evolved’ population increased significantly from 0.64 h⁻¹ to 1.196 h⁻¹ (Tukey’s HSD, *p-value* < 0.001), though still below the WT rate (1.3 h⁻¹). Similarly, the ‘evolved-WT’ population showed a growth rate improvement from 1.3 h⁻¹ to 1.6 h⁻¹ (Tukey’s HSD, *p-value* < 0.001) (**Fig. 1B, Table S1**). To identify the accumulated mutations, we isolated three random variants from both the ‘evolved’ and ‘evolved-WT’ populations from the last serial passage (generation 320), validated that their growth rates closely match the average growth rates of their respective populations (**Fig. S2**), and performed whole genome sequencing (**Table 1**). There were no shared mutations between the two sets, indicating that the *metK* replacement profoundly reshaped the mutational fitness landscape of the ‘unevolved’ strain. All ‘evolved-WT’ clones carried a known 82 bp deletion in the *rph-pyrE* operon, restoring pyrimidine synthesis^17, 51, 52^, whereas all ‘evolved’ clones shared two fixed mutations: a *metK* promoter mutation and a nonsense mutation in the pleiotropic *pntA*, inactivating transhydrogenase and shifting NADH/NADPH balance^35, 53^. Other mutations, such as in pleiotropic genes *rpoB* (encoding RNA polymerase subunit beta^46^) and *prfC* (encoding peptide chain release factor 3^47^), appeared sporadically and have been seen in other lab-evolution studies (referenced in **Table 1**), suggesting potential adaptive roles even though they did not reach fixation.

**Table 1.** Mutations found in isolated variants.

| Gene | Mutation | 'evolved' population<br>*isolated variants* |  |  | 'evolved-WT' population<br>*isolated variants* |  |  | Ref. |
| --- | --- | --- | --- | --- | --- | --- | --- | --- |
|  |  | *v1* | *v2* | *v3* | *v1* | *v2* | *v3* |  |
| *pntA* | Nonsense<br>CAA→TAA (Q322STP) | ✓ | ✓ | ✓ |  |  |  | 35, 53-56 |
| *metK* | Intergenic (-8) T→A | ✓ | ✓ | ✓ |  |  |  | This work |
| *prfC* | Missense<br>GAT→AAT (D222N) | ✓ |  |  |  |  |  | 52, 57 |
| *fes* | Intergenic (-89) G→T | ✓ |  |  |  |  |  | 56, 58 |
| *yagE* | Intergenic (-70) ΔA | ✓ |  |  |  |  |  |  |
| *rpoB* | Missense<br>CAC→TAC (H526Y) |  | ✓ |  |  |  |  | 17, 46, 52, 54 |
| *flgB* | In frame deletion<br>ΔTAACAGCCT<br>(1131349) |  |  | ✓ |  |  |  | 59 |
| *sufS* | Frameshift deletion ΔT<br>(1759368) |  |  | ✓ |  |  |  | 55, 60 |
| *rph-pyrE* | Δ82 bp (3815848) |  |  |  | ✓ | ✓ | ✓ | 17, 51, 52 |
| *flhD* | Frameshift insertion<br>C (1978164) →CGGTG |  |  |  | ✓ | ✓ |  | 61, 62 |
| *insAB1* | Gene duplication<br>(1978500-1979250) |  |  |  | ✓ | ✓ |  |  |
| *paoC* | Missense<br>GAT→GTT (D13V) |  |  |  |  | ✓ |  |  |
| *cadA* | Synonymous<br>TCC→TCT (S331) |  |  |  |  |  | ✓ |  |

For further analysis, we have arbitrarily chosen ‘evolved *v1*’ and ‘evolved-WT *v1*’ variants (from now on, ‘evolved’ and ‘evolved-WT’ strains, respectively). We found that evolution significantly increased the intracellular UuMAT abundance in the ‘evolved’ strain by ∼1.4-fold (Pairwise *t*-test with Bonferroni adjustment, *p-value* < 0.001) (**Fig. S1B**) and alleviated metabolic imbalance by significantly raising SAM (Tukey’s HSD, *p-value* < 0.001) and reducing methionine levels (*p-value* < 0.001) (**Fig. S1A**). In principle, a recurrent mutation in the *metK* promoter might solely account for these changes by boosting UuMAT expression, potentially rendering other accumulated mutations inconsequential to the observed phenotypic and growth improvements. To isolate the effects of the *metK* promoter mutation, we reintroduced that single mutation into the ancestral background (’mut’ strain). Although ‘mut’ matched the ‘evolved’ strain in UuMAT abundance (**Fig. S1B**), its growth rate remained slightly lower (1.173 h^-^¹ vs. 1.196 h^-^¹), whereas its lag phase grew significantly longer (62 min vs. 53 min; One-way ANOVA across strains: F_4,5_ = 199.3, *p-value* = 1.07·10^-5^, followed by Tukey’s HSD, *p-value* < 0.05) (**Fig. 1B**, **Table S1**). Moreover, ‘mut’ accumulated significantly more SAM than the evolved strain (Tukey’s HSD, *p-value* < 0.001) despite indistinguishable methionine levels, yielding a significantly higher SAM/methionine ratio (One-way ANOVA across strains: F4,5 = 463.4, *p-value* < 2·10^-16^, followed by Tukey’s HSD, *p-value* < 0.001) (**Fig. S1A**).

Thus, although the mutation in the *metK* promoter clearly demonstrates a strong adaptive value and has likely reached fixation through a selective sweep during the early stages of evolution, additional mutation(s) that have accumulated in the ‘evolved’ variant (**Table 1**) further modified the metabolic phenotype and enhanced the fitness of the ‘evolved’ strain.

In the ‘evolved-WT’ strain, EcMAT abundance and SAM levels both rose significantly (EcMAT, *p-value* = 0.0194, *t*-test; SAM, *p-value* = 0.003, Tukey’s HSD), but methionine levels also increased (Tukey’s HSD, *p-value* = 0.03), keeping the SAM/methionine ratio at wild-type levels (**Fig. S1**). Because none of the ‘evolved-WT’ mutations overlap with those in the ‘evolved’ strain, the shifts in MAT and SAM likely reflect broader metabolic and regulatory changes driven by its distinct mutational trajectory (**Table 1**). Collectively, these results underscore that the mechanisms and outcomes of metabolic adaptation depend strongly on a strain’s initial metabolic state.

### Strains show distinct metabolic states, separating evolution and genome editing

Bacterial populations propagated under seemingly identical laboratory conditions often display significant variability in fitness across replicates, even when fitness is measured with high precision^33^. This variability is further intensified when measurements are taken on different days, a phenomenon known as the ‘batch effect’^33^. Fitness variability is primarily attributed to physiological sensitivity to subtle environmental perturbations that inevitably exist between replicates in laboratory settings, due to minor temperature gradients in the incubator, slight differences in vessel shape, and variations in shaking speed. Given the highly dynamic nature of the metabolome and its hyper-responsiveness to environmental conditions^23^, miniscule environmental changes are likely to induce even greater variability in metabolite levels and, potentially, alter metabolite correlation patterns.

To account for the metabolic variability and identify metabolite correlation patterns that withstand subtle environmental perturbations, we implemented the following strategy. First, we increased the number of replicates in a single batch experiment from the standard ten replicates^24^ to nineteen. Second, we conducted two such batch experiments on different dates, resulting in a total of 38 replicates per strain. This strategy was applied to the WT, ‘evolved-WT’, ‘unevolved’, ‘evolved’, and ‘mut’ strains. Each replicate was grown to a mid-exponential phase and subjected to polar metabolite extraction and LC-MS metabolic profiling under positive ionization mode (**Materials and Methods)**. The processing of raw chromatograms produced data files containing the relative abundances of all individual untargeted metabolic features. In each replicate, we identified 2,118 identical metabolic features (**Fig. 1C, Fig. S3, Supplementary Data**).

We performed Principal Component Analysis (PCA) on the metabolite abundance data separately for each batch and found a clear strain-specific clustering, although differences between the batches were also apparent. For example, the ‘evolved-WT’ and ‘evolved’ strains were more distinctly separated along both PC1 and PC2 in batch 1 compared to batch 2 (**Fig. S4**). However, when we combined the data from both batches into a single dataset, the PCA showed a sharp, strain-specific group separation, with strong clustering of replicate data points from both batches **(Fig. 1D**). The largest shift along PC1 was caused by the orthologous *metK* replacement (from the WT to the ‘unevolved’ strains). Evolutionary adaptation (’evolved’ strain), or the introduction of a single mutation in the *metK* promoter (’mut’ strain) shifted the data points back towards the WT strain. Notably, the better growing ‘evolved’ strain moved closer to the WT than the ‘mut’ strain, suggesting that PC1 reports on metabolic optimality. Conversely, the evolutionary adaptation of the ‘evolved-WT’ strain, which accumulated a completely different set of mutations compared to the ‘evolved’ strain, pushed the data points away from the WT along PC2, indicating that PC2 likely captures diversity among optimal metabolic states (**Fig. 1D**). Taken together, the PCA analysis demonstrates that each strain represents a distinct metabolic state, with differences reflecting their evolutionary relationships. Combining the metabolite abundance data from two batch experiments further strengthens these distinctions.

### Metabolite correlation networks reveal strain-specific rewiring while preserving a core of robust correlations

Having established that each strain is characterized by a unique metabolite abundance pattern, we turned to constructing the metabolite correlation networks. To this end, we converted the metabolite abundances into correlation matrices using Pearson’s correlations. A separate correlation matrix was constructed for each batch (**Fig. 1C**). To reduce the contribution of non-specific transitive correlations^25^, the obtained matrices were filtered by the Probabilistic Context Likelihood of Relatedness (PCLRC) algorithm^63^ (see **Materials and Methods**). Finally, to construct a network comprised of correlations that withstand subtle environmental perturbations, we retained only the correlations observed in networks of both batches, resulting in a single overlap network per strain (**Fig. 1C**). In all strains, the Jaccard correlations between both batches were rather small (0.126-0.156, **Fig. 1E**, diagonal), indicating the dramatic influence of environmental conditions on metabolite correlation patterns.

Jaccard similarity analysis across overlap networks of individual strains also revealed low levels of shared links ranging from 0.098 (between the WT and ‘unevolved’ networks) to 0.19 (between the ‘evolved’ and ‘mut’ networks) (**Fig. 1E**, off-diagonal). The unsupervised hierarchical clustering analysis performed on the Jaccard similarity values brought together the WT and ‘evolved-WT’ networks and the ‘evolved’ and ‘mut’ networks, while placing the ‘unevolved’ network at an equal distance from the rest of the networks (**Fig. 1E**), thus capturing the hierarchy of network separation observed in PCA (**Fig. 1D)**.

To estimate the significance of Jaccard similarity analysis performed across the overlap networks, we shuffled each empirical overlap network 1,000 times to create random networks, while keeping the number of links per node. The observed Jaccard values were significantly different (*p-value* << 10^-4^, one-sample *z*-test) (**Fig. S5**), indicating that the empirical networks are more similar to each other than expected by chance. Therefore, despite the massive rewiring of correlation networks caused by genetic changes, there remains a common robust core of correlations that withstood genetic perturbations. For instance, the ‘evolved-WT’ strain retained 36% of the correlations found in the ancestral WT strain, while the ‘evolved’ strain preserved 25% of the correlations detected in the ancestral ‘unevolved’ strain (**Fig. S6**). The ‘mut’ strain, carrying the single mutation in the *metK* promoter, and the ‘evolved’ strain carrying, in addition to the *metK* promoter mutation, four off-target mutations (**Table 1**), shared only 34.3% and 29.9% of the correlations, respectively, indicating the dramatic effect of the off-target mutation(s) on network structure. A small number (2,772) of correlations shared between the ‘evolved’ and ‘evolved-WT’ networks (9% and 8.5% of the correlations in the ‘evolved’ and ‘evolved-WT’ strains, respectively) were not found in the WT network, indicating that although the ‘evolved’ and ‘evolved-WT’ strains did not accumulate common mutations, they nonetheless shared some of the metabolic adaptations, possibly because both have evolved under identical conditions. The number of core correlations shared by all five strains amounted to 2,477, representing only 4.7% of the correlations found in the WT network (**Fig. S6**).

### Adaptive evolution eliminates links and removes hubs

To evaluate the impact of mutations on the structure of metabolite correlation networks, we considered separately positive and negative correlations. While the full scope of mechanisms affecting the metabolite correlation sign remains unclear, computational modeling suggests that *positive* correlations only occur when two metabolites approach chemical equilibrium, where their ratio converges to the equilibrium constant. *Positive or negative* correlations may arise (i) under asymmetric control, when the concentration of a dominant metabolite that controls the concentration of two other metabolites fluctuates; or (ii) under regulation by a common transcription factor. *Negative* correlations only are expected (i) between metabolites involved in conserved moiety production/degradation cycles (due to mass conservation); or (ii) between a substrate and a product of an enzymatic reaction catalyzed by an enzyme with a highly variable expression ^23–26, 64^. While there are no apparent mechanistic reasons to anticipate the high prevalence of any of these mechanisms, we found that positive correlations dominate the networks. They account for 78.7% to 88.6% of all correlations, suggesting that mechanisms underlying positive correlations constitute the main driver of concentration interdependence between metabolites (**Table 2**). Similar levels of positive versus negative correlations in metabolite and transcriptome correlation networks were reported in previous studies^65–67^.

**Table 2.**
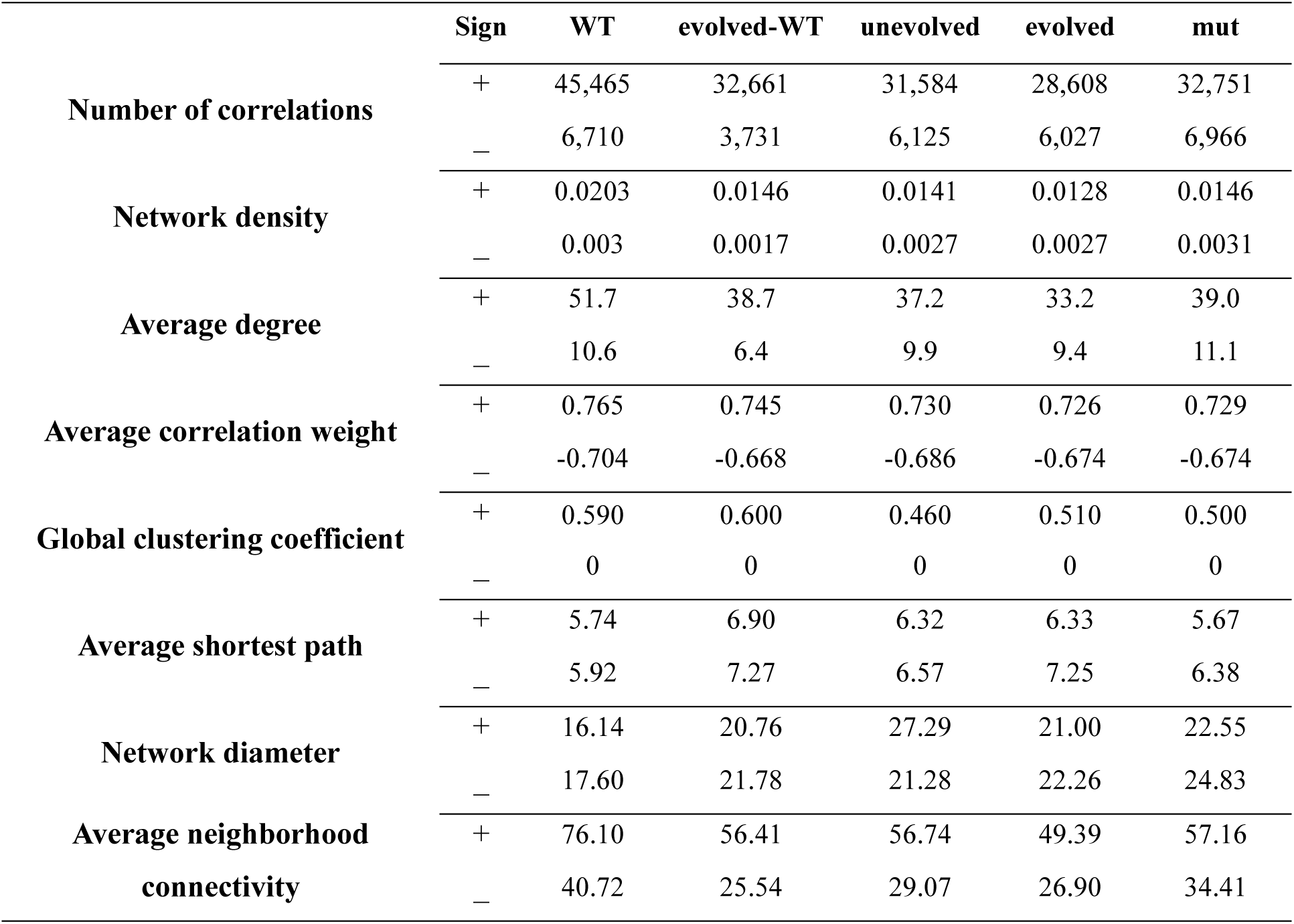
Global network properties.

|  | Sign | WT | evolved-WT | unevolved | evolved | mut |
| --- | --- | --- | --- | --- | --- | --- |
| Number of correlations | + | 45,465 | 32,661 | 31,584 | 28,608 | 32,751 |
|  | – | 6,710 | 3,731 | 6,125 | 6,027 | 6,966 |
| Network density | + | 0.0203 | 0.0146 | 0.0141 | 0.0128 | 0.0146 |
|  | – | 0.003 | 0.0017 | 0.0027 | 0.0027 | 0.0031 |
| Average degree | + | 51.7 | 38.7 | 37.2 | 33.2 | 39.0 |
|  | – | 10.6 | 6.4 | 9.9 | 9.4 | 11.1 |
| Average correlation weight | + | 0.765 | 0.745 | 0.730 | 0.726 | 0.729 |
|  | – | -0.704 | -0.668 | -0.686 | -0.674 | -0.674 |
| Global clustering coefficient | + | 0.590 | 0.600 | 0.460 | 0.510 | 0.500 |
|  | – | 0 | 0 | 0 | 0 | 0 |
| Average shortest path | + | 5.74 | 6.90 | 6.32 | 6.33 | 5.67 |
|  | – | 5.92 | 7.27 | 6.57 | 7.25 | 6.38 |
| Network diameter | + | 16.14 | 20.76 | 27.29 | 21.00 | 22.55 |
|  | – | 17.60 | 21.78 | 21.28 | 22.26 | 24.83 |
| Average neighborhood connectivity | + | 76.10 | 56.41 | 56.74 | 49.39 | 57.16 |
|  | – | 40.72 | 25.54 | 29.07 | 26.90 | 34.41 |

We began network analysis by calculating network density, a global property that quantifies the ratio of observed to possible links. Every genetic perturbation, whether engineering (WT → ‘unevolved’) or evolution (WT → ‘evolved-WT’ and ‘unevolved’ → ‘evolved’), consistently lowered network density (**Table 2**). WT → ‘unevolved’ and WT → ‘evolved-WT’ transitions each removed about 30% of positive links, while negative links fell by 45 % only in ‘evolved-WT’, highlighting the difference in the metabolic states between the ‘unevolved’ and ‘evolved-WT’ strains (**Table 2**). In contrast, the ‘unevolved’ → ‘evolved’ transition cut positive links by 10 % and negative links by less than 1 %. Link counts were also lower in the ‘evolved’ than in the ‘mut’ strain, indicating the effects of additional off-target mutation(s). Link loss coincided with weaker correlation weights: both the ‘evolved-WT’ and ‘unevolved’ networks showed significant downward shifts in positive and negative weights (*p-value* < 2·10^-16^, Wilcoxon rank sum test with BH-fdr correction), with average weight of positive correlations dropping from 0.765 (WT) to 0.745 (’evolved-WT’). The weight reduction in the ‘unevolved’ → ‘evolved’ transition was smaller, but still significant (*p-value* < 2·10^-15^). The average weight of positive correlations was also slightly lower in the ‘evolved’ than in ‘mut’ network (*p-value* < 3·10^-8^), while negative weights remained unchanged (*p-value =* 0.66). (**Table 2**, **Fig. S7**).

These findings indicate that rewiring in the ‘evolved-WT’, ‘unevolved’, and ‘evolved’ networks produced generally weaker correlations compared to those seen in the corresponding ancestral networks. Furthermore, the average weights of shared positive and negative correlations in the WT/’evolved-WT’, WT/’unevolved’, and ‘unevolved’/’evolved’ ancestral/derived pairs had higher mean weights than network-wide averages (*p-value* < 0.01), indicating that stronger metabolite correlations are more resilient to genetic perturbations. A similar conclusion is held when shared correlations between the ‘evolved’ and ‘mut’ networks are compared (**Table S2, Fig. S8**).

To better understand how link loss and weight reduction shaped the properties of individual networks, we compared the distributions of node degree (number of links) and found that density loss stemmed mainly from removal of highly connected nodes (hubs) (**Fig. 2A**). This is further evidenced from the direct comparisons of the degree distributions of the top 5% degree nodes, which show a significant decrease in both the positive and negative node degree in the ‘evolved-WT’ network (*t*-test *p-values* < 2·10^-16^), and in the positive node degree of the ‘unevolved’ and ‘evolved’ networks (*p-value* < 2·10^-5^) (**Fig. 2B**). The top 5% degree nodes were also significantly lower in the ‘evolved’ network than in the ‘mut’ network (*p-value* = 0.017 and *p-value* < 0.001, for positive and negative correlations, respectively) indicating that the accumulation of the off-target mutation(s) in the ‘evolved’ strain on top of the *metK* promoter mutation in the ‘mut’ strain was associated with a loss of network hubs (**Fig. 2B**).

**Fig. 2.**
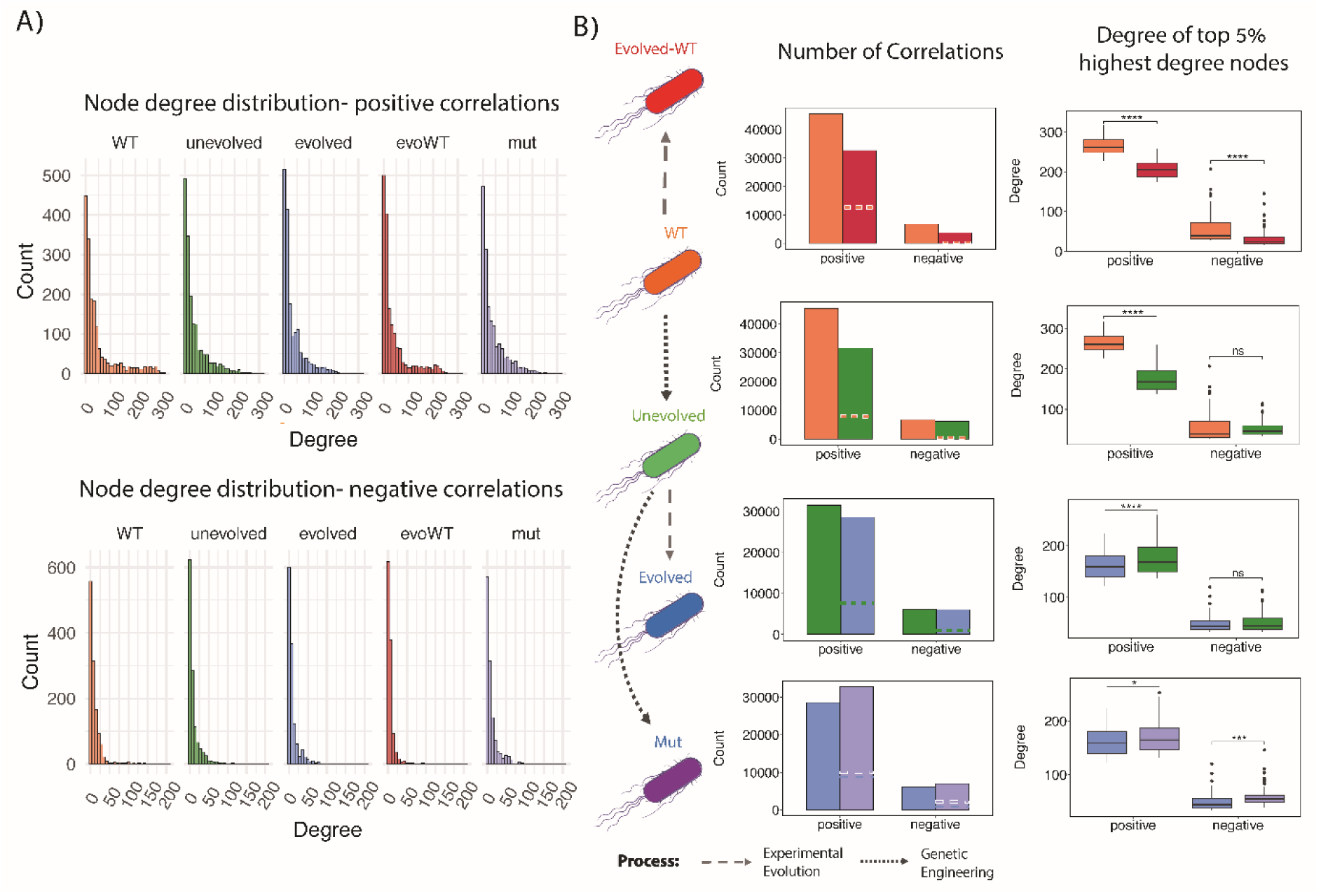
Analysis of node degree. **A.** Network degree distributions. *Upper panel,* positive correlations; *lower panel,* negative correlations. **B.** *Left panel*, color coding of network strains; *middle panel*, total number of positive and negative correlations in each network. Dashed lines indicate the number of common correlations between the two compared networks. *Right panel,* comparisons of node degree between the top 5% nodes with the highest degree in each network. **** *p-value* <10^-4^, *** *p-value* <10^-3^ (Two Sample *t*-test); ns – nonsignificant.

The reduction in network hubs is also expected to influence the node neighborhood connectivity (NC), which quantifies the average number of links of the neighboring nodes. Indeed, we found that hub loss reduced NC: top 5 % NC nodes and average NC fell across all derived networks (for both positive and negative correlations) (**Table 2, Fig. S9**). The steepest decline, about 25 % in mean positive NC (*t*-test, *p-values* < 2·10^-16^), occurred in the ‘evolved-WT’ and ‘unevolved’ networks, whereas the ‘evolved’ network showed a two-fold smaller drop (*p-value* < 2·10^-10^). Average NC and top 5 % NC were also markedly lower in the ‘evolved’ than in ‘mut’ network (*p-values* < 10^-8^) (**Table 2**, **Fig. S9**).

Altogether, these findings demonstrate that both the replacement of the *metK* gene to generate the maladapted ‘unevolved’ strain and adaptive evolution to produce the ‘evolved-WT’ and ‘unevolved’ strains led to an elimination of a substantial fraction of metabolite correlations and to a reduction in the number of highly correlated metabolites (node hubs), potentially affecting the overall connectivity of the metabolite correlation networks. In addition, the strength of the newly formed metabolite correlations (link weights) within these networks was significantly weaker compared to the correlation weights in the WT network, indicating that the cohesiveness of these networks (a propensity to form closely-knit groups) was also negatively affected. The substantial differences in the density, average node degree, neighborhood connectivity, and correlation weights between the ‘evolved’ and ‘mut’ networks further indicate that the off-target mutation(s) present in the ‘evolved’ strains in addition to the *metK* promoter mutation are responsible for a profound restructuring of the metabolite correlation network.

### Evolution reshapes the ‘small-world’ properties of metabolite correlation networks

Although changes in the global properties of the ‘evolved-WT’, ‘unevolved’, and ‘evolved’ networks overall followed a similar trend, the magnitude of changes accompanying the WT → ‘evolved-WT’ and WT → ‘evolved-WT’ transitions was always much more pronounced than this observed in the ‘unevolved’ → ‘evolved’ transition. For instance, as demonstrated above, the ‘evolved’ network lost three times fewer positive links and exhibited a two-fold smaller reduction in the average neighborhood connectivity than the ‘evolved-WT’ and ‘unevolved’ networks (**Table 2**). Seeking an explanation for this phenomenon, we noticed that a particular set of values, which are key to the networks’ ‘small-world’ properties, diverged from the general trend seen in the other global properties across all three networks. A key characteristic of ‘small-world’ networks is a balance between functional segregation of nodes into local clusters (modules) and the integration of information flow between these clusters^68^. The dense local connectivity within ‘small-world’ networks ensures efficient communication within clusters, whereas the sparse connections between clusters support fast integration of information at the global network level. The magnitude of network segregation into clusters can be quantified with the Global Clustering Coefficient (GCC), a parameter derived from the clustering coefficient of individual nodes (**Materials and Methods**). The latter measures the extent to which nodes connected to a focal node are also connected to each other, forming a triangle^69^. The global clustering coefficient (GCC) assesses the overall level of cliquishness or local grouping within a network with values ranging from 0 to 1. A value of 0 indicates no clustering, while a value of 1 means every connected trio of nodes forms a triangle^68^. Network integration, in turn, is quantified by the average shortest path length (*L*) across all node pairs. Unlike regular networks, characterized by a high GCC and a large *L*, or random networks that display a low GCC but a small *L*, ‘small-world’ networks are defined by a relatively high GCC and a relatively small *L*. Using these assessments, we determined that the metabolite correlation network of the WT strain exhibits ‘small-world’ properties (see **Supplementary Note I**). Many biological networks, including metabolic, protein-protein interaction, and brain networks, are also known to display ‘small-world’ characteristics^69–71^.

As the WT → ‘evolved-WT’, WT → ‘unevolved’, and ‘unevolved’ → ‘evolved’ network transitions took place, they underwent distinct patterns of changes in their GCC and average shortest path length values. The ‘evolved-WT’ network preserved its GCC, suggesting that the level of connectivity within its densely interconnected clusters remained largely stable, despite losing about 30% of positive and 45% of negative links and undergoing extensive rewiring (**Table 2, Fig. S6**). This observation is further supported by a correlation analysis between node clustering coefficients and node degrees, which shows that in the ‘evolved-WT’ network, nodes with high degree retained clustering coefficients, in similarity to those in the ancestral WT network (**Fig. S10**). Unlike the Global Clustering Coefficient (GCC), the average shortest path length in the ‘evolved-WT’ network increased for both positive and negative correlations, rising from 5.74 to 6.9 and from 5.92 to 7.27, respectively (**Table 2**). Similarly, the network diameter, defined as the longest shortest path between any two nodes, also increased from 16.14 to 20.76 for positive and from 17.6 to 21.78 for negative correlations, suggesting a decrease in the efficiency of information flow between clusters (**Table 2**).

In contrast to the ‘evolved-WT’ network, the ‘unevolved’ network displayed a substantial reduction in its GCC, which declined from 0.59 to 0.46, suggesting that the node hubs lost during the transformation from the WT network were integral to highly interconnected local clusters (**Table 2**). Indeed, as can be seen from the analysis of correlations between node clustering coefficient and node degree, the high degree nodes in the ‘unevolved’ network have substantially lower clustering coefficients compared to their counterparts in the WT network (**Fig. S10**). In similarity to the ‘evolved-WT’ network, the average shortest path length in the ‘unevolved’ network has also increased, albeit to a lesser extent, from 5.74 to 6.32 for positive correlations and from 5.92 to 6.57 for negative ones (**Table 2**). Based on these findings, we concluded that WT → ‘unevolved’ transformation was accompanied by a deterioration in the ‘small-world’ properties of the ‘unevolved’ network, likely due to the removal of node hubs that led to a partial disintegration of highly interconnected local groups.

In contrast to the ‘evolved-WT’ network, the adaptation of the ‘evolved’ strain has started from the low-fitness ‘unevolved’ strain with low network density and disintegrated ‘small-world’ properties. Evolution led to a strengthening of the ‘small-world’ properties, as evidenced by the increase in the GCC from 0.46 (in the ‘unevolved’ network) to 0.51, the preservation of the average shortest path for positive correlations, and a reduction in network diameter from 27.29 to 21 for positive correlations (**Table 2**). These changes in the ‘evolved’ network can be attributed to the improved local connectivity between nodes within network clusters through the addition of positive correlations, a process that has taken place despite an overall reduction in the number of positive links (**Table 2**). The necessity to restore the local connectivity within clusters of the ‘evolved’ network can explain why the drop in most global network parameters (network density, average link weight, average neighborhood connectivity) seen in the WT → ‘evolved-WT’ and WT → ‘unevolved’ network transformations was much less pronounced in the ‘evolved’ strain.

Importantly, the GCC of the ‘evolved’ network was also slightly higher than that of the ‘mut’ network (0.510 vs 0.500), suggesting that the off-target mutation(s) that have accumulated in the ‘evolved’ strains contributed to the improvement of the local node connectivity within network clusters on top of the improvement achieved by the *metK* promoter mutation (GCC of 0.50 in the ‘mut’ network vs GCC of 0.46 in the ‘unevolved’ network) (**Table 2**). The off-target mutation(s) also shortened the network diameter of the ‘evolved’ network for both positive and negative correlation (from 22.55 in the ‘mut’ network to 21.00 for positive, and from 24.83 in the ‘mut’ network to 22.26 for negative correlations) (**Table 2**). This change, which reports on the removal of the longest shortest path between two nodes in a network, however, was not accompanied by a reduction in the average shortest path of the ‘evolved’ network that went up from 5.67 in the ‘mut’ network to 6.33 (for positive correlations), and from 6.38 to 7.25 for the negative ones (**Table 2**). This inconsistency probably suggests that the global improvement in the information flow between network clusters achieved by the *metK* promoter mutation (as reported by the average shortest path) does not automatically ensure better fitness. Indeed, as stated above, the evolutionary adaptation from the WT to the ‘evolved-WT’ strains was also marked by a reduction in the average shortest path.

### Changes in network node roles and properties reveal adaptive phenotypes of mutations

So far, our global-level analysis has identified mutation-induced changes in the ‘small-world’ properties of the networks without providing detailed information about changes in the local network structure or the properties of individual nodes. To uncover these details, we first performed a meso-scale network analysis. For each strain, we constructed bilayer networks separating positive and negative correlations into distinct layers and partitioned these networks into modules using the Infomap algorithm (see **Materials and Methods**). Although the overall distribution of module sizes appeared similar across all networks, key differences emerged in the absolute sizes of large and medium modules, the composition of correlation signs within modules (entirely positive, entirely negative, or mixed), and the average node properties within modules, namely, clustering coefficient, degree, and betweenness (**Fig. 3A, Table S3**). For instance, while the largest module in the WT network contained 499 nodes, the largest module in the ‘evolved-WT’ network comprised only 402 nodes. Similarly, the ‘unevolved’ → ‘evolved’ transition involved a 34% reduction in the size of the largest module (from 521 to 344 nodes). This reduction was accompanied by significant increases in average clustering coefficient (from 0.50 to 0.53, Wilcoxon rank sum test with continuity correction, *p-value* = 0.002), average betweenness (from 0.0017 to 0.0019, *p-value* = 0. 048), and average node degree (from 82.2 to 86.6, p-value < 2.2·10^-16^) (the largest modules in both the ‘unevolved’ and ‘evolved’ networks consisted entirely of positively correlated nodes) (**Fig. 3A, Table S3**). Furthermore, the four largest modules in the ‘evolved’ network were approximately 13–28% smaller than their counterparts in the ‘mut’ network, underscoring the impact of off-target mutation(s) on meso-scale network structure (**Fig. 3A, Table S3).**

**Fig. 3.**
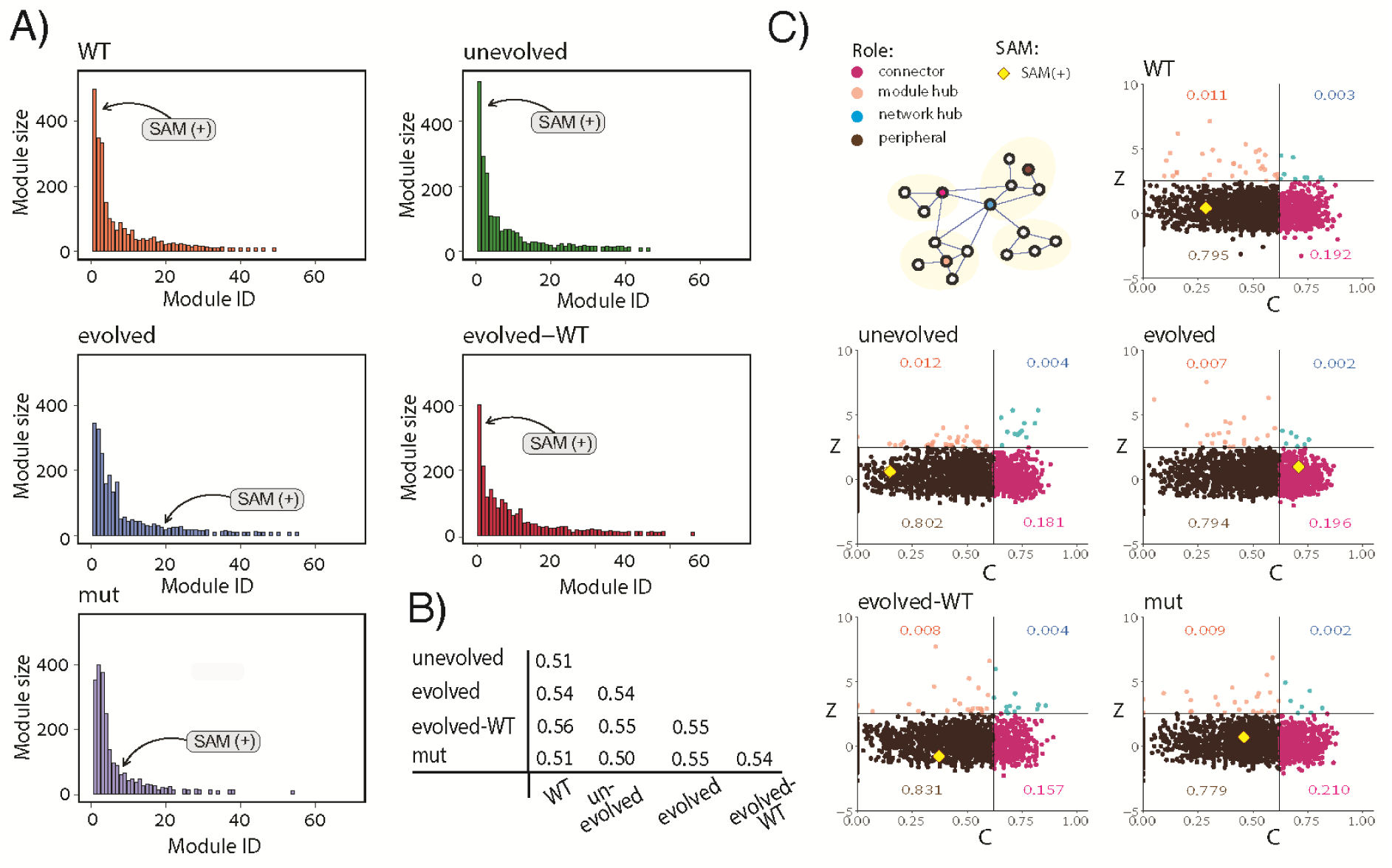
Change in node roles upon genetic engineering and adaptive evolution. **A**. Distribution of network module sizes (number of nodes). Only modules with 10 nodes or more are shown. Arrows point to modules that include SAM (only SAM nodes involved in positive correlations are shown). **B.** Normalized Mutual Information (NMI) analysis of the module composition among networks. **C.** Node roles. Each dot represents a node with an assigned role according to its connectivity within (*Z*) and between (*C*) modules. Color-coded numbers indicate the relative proportion of nodes with the assigned role. Nodes representing SAM involved in positive correlations are depicted as yellow diamonds.

Together, these results suggest that WT → ‘evolved-WT’ and ‘unevolved’ → ‘evolved’ adaptive evolutions led to a fragmentation of the networks into smaller, more cohesive modules, while preserving or even enhancing intramodular connectivity. These meso-scale structural changes offer an explanation for the preservation or increase in network global clustering coefficients, despite an overall reduction in the number of network links in evolution (**Table 2**).

To further assess the structural differences between networks, we performed a Normalized Mutual Information (NMI) analysis of modules’ composition (see **Materials and Methods**). This analysis revealed only moderate similarity between network modules, with pairwise comparisons across all networks ranging from 0.5 to 0.56 (**Fig. 3B**), hinting that properties of individual nodes may have changed substantially between networks. To probe this hypothesis, we identified and characterized the properties of the node representing SAM, the product of MAT-catalyzed reaction. SAM is allocated to the largest module in the WT, ‘unevolved’, and ‘evolved-WT’ networks (when only positive interactions are considered). Strikingly, the ‘unevolved’ to ‘evolved’ strain evolution resulted in the relocation of SAM into a much smaller node (**Fig. 3A**). SAM was also found in a bigger module in the ‘mut’ network, further demonstrating the unique impact of off-target mutation(s) on the ‘evolved’ network’s structure and function (**Fig. 3A**).

Encouraged by this finding, we aimed to develop a comprehensive strategy to identify all the nodes potentially important for metabolic adaptation. Following the previously developed method^72^, we categorized all network nodes based on their roles, defined by their within-module degree (*Z*), which quantifies a node’s connectivity within its module, and the participation coefficient (*C*), which measures the distribution of a node’s connections across different modules (see **Materials and Methods**). This strategy revealed that across all networks, peripheral nodes that form only weak connections within their respective modules were the most abundant, comprising between 79.4% and 83.1% of all nodes (**Fig. 3D**). Connectors, which have high connectivity to nodes outside their modules, constituted the second most abundant group, accounting for 15.7% to 21% of the nodes (**Fig. 3D**). Module and network hubs, representing nodes with significant within-module or network-wide connections, respectively, were the least abundant, collectively ranging from 9% to 16% of all nodes (**Fig. 3D**). The role of the node representing SAM shifted from peripheral to connector between the ‘unevolved’ and ‘evolved’ networks (for positive correlations). Notably, SAM assumes the connector role exclusively in the ‘evolved’ network, again underscoring the substantial impact of the off-target mutation(s). This transition to the connector role was accompanied by a seven-fold decrease in node degree, an almost two-fold decrease in clustering coefficient, and a five-fold reduction in neighborhood connectivity, compared to the ‘unevolved’ network (for positive correlations) (**Fig. 4A-C**). In fact, these parameters reached their lowest levels in the ‘evolved’ network relative to all other networks. In stark contrast, the betweenness centrality of SAM, which is determined by the number of shortest paths passing through the node, increased by at least 10-fold in the ‘evolved’ network compared to all other networks **(Fig. 4D).**

**Figure 4.**
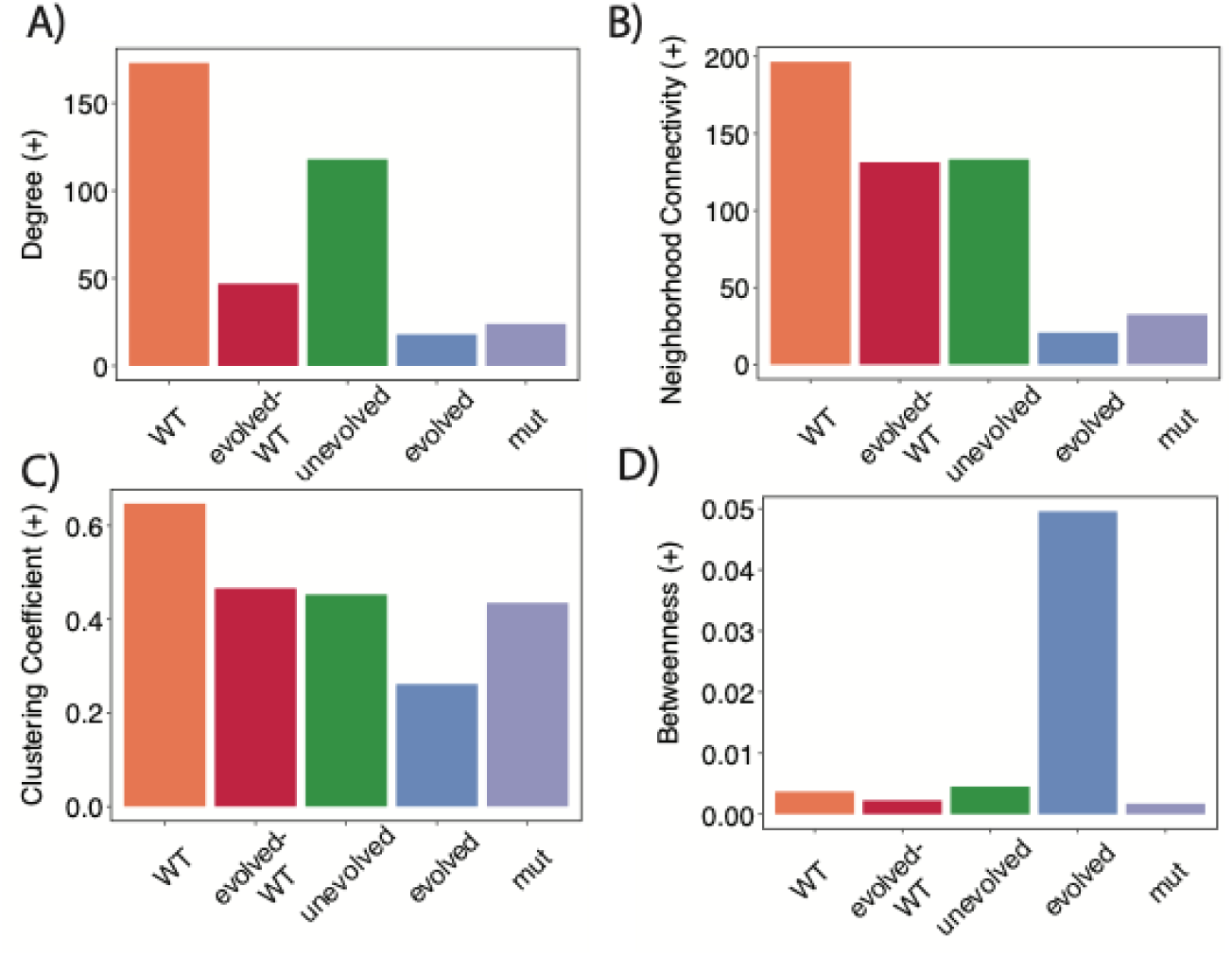
SAM node properties in the correlation networks (for positive correlations). **A.** Node degree**. B.** Neighborhood connectivity. **C**. Clustering coefficient. **D**. Betweenness centrality.

Collectively, these findings suggest that the transition from the ‘unevolved’ to the ‘evolved’ strain not only relocated SAM from the largest to a smaller module but also reduced its centrality within that module by severing interactions with tightly connected node groups. At the same time, SAM’s role in linking different modules became more pronounced in the ‘evolved’ network, potentially indicating improved coordination of SAM metabolism. Notably, the clustering coefficient and betweenness centrality of the SAM node in the ‘mut’ strain more closely resembled those in the ‘unevolved’ network than in the ‘evolved’ one (**Fig. 4C, D**).

### Targeted metabolomics of key nodes provides a novel path towards mechanistic exploration of mutation effects

The informative changes observed in the SAM node during the transition from the ‘unevolved’ to the ‘evolved’ strain suggest that identifying other nodes exhibiting similar shifts in role, connectivity, and centrality could generate testable hypotheses for the mechanisms underlying metabolic adaptation. To pursue this, we analyzed the ‘evolved’ network and identified: 71 connector nodes that shifted from peripheral roles in the ‘unevolved’ network but remained peripheral in all other networks; 111 nodes with at least 40% reduced clustering coefficients compared to the ‘unevolved’ network; and 71 nodes with at least 5-fold increased betweenness centrality relative to the ‘unevolved’ network (**Table S4**). Using deep neural network–based predictions of high-resolution fragmentation spectra from LC-MS/MS data^73–76^, we successfully assigned putative identities to 18 of these nodes (**Table S5**; see **Materials and Methods**). Among these were SAM and 5-methylthioadenosine (MTA), whose identities were validated using authentic standards (**Materials and Methods**).

SAM is part of a tightly interconnected network of metabolic pathways involved in the synthesis of purines, methionine, and polyamines^50, 77, 78^. We found that five of the predicted metabolites (L-glutamine, SAICAR, 5,10-methenyltetrahydrofolate, xanthosine, and S-hydroxymethyl glutathione) participate in de novo or salvage purine biosynthesis, two (L-threonine and L-homoserine phosphate) are involved in regulation of de novo methionine synthesis, and three (palmitoylputrescine, N-acetylputrescine, and methylthioadenosine) are linked to polyamine biosynthesis. Three additional metabolites, enterobactin, cyclic AMP (cAMP), and γ-glutamylcysteine, play global roles in *E. coli* metabolism, physiology, and growth (**Table S5** and **Supplementary Note II**).

Overall, the analysis indicates that targeted metabolomics of nodes whose network properties shift under adaptive pressure can yield potentially relevant candidates for further mechanistic exploration of metabolic adaptation during evolution.

## Discussion

In this work, we developed an experimental and analytical framework to examine how mutations alter metabolite correlation networks. We demonstrate that this approach provides a practical tool for exploring genotype–phenotype–fitness relationships underlying metabolic adaptation in bacteria. Furthermore, we propose that targeted metabolomics of key network nodes whose roles shifted in response to adaptive mutations can generate testable hypotheses to guide further investigation into the metabolic mechanisms of adaptation.

Unlike topological metabolic networks where nodes represent metabolites and edges denote biochemical transformations mediated by enzymes, metabolite correlation networks do not necessarily capture direct biochemical relationships or causality. Instead, two metabolites might be highly correlated due to shared regulation or association rather than being directly connected in a metabolic pathway. The correlation between two metabolites is a *systemic* property influenced by the entire metabolic network, not just the direct interaction between those two metabolites. Thus, the observed pattern of correlations in a metabolite correlation network is considered a global ‘fingerprint’ of the physiological state of the system under given conditions. The structure and function of the metabolite correlation networks are, therefore, *emergent properties* arising from the cumulative effects of biochemical reactions and regulatory mechanisms. These networks dynamically adjust to genetic and environmental changes, reflecting the newly established physiological state. Consequently, changes in network properties in response to genetic perturbations constitute complex phenotypes that link adaptive mutations to bacterial fitness. The detected changes, however, do not provide a direct mechanistic explanation of metabolic adaptation.

The most advanced tool for understanding the mechanisms of adaptation to inefficient enzymes is Metabolic Flux Analysis (MFA)^35, 79^. MFA relies on tracing the incorporation of labeled substrates into downstream metabolites to quantify changes in reaction rates (fluxes) in response to mutations^16^. This approach crucially depends on the detailed knowledge of the metabolic network and on detection of metabolites in the affected pathways. Although it is generally assumed that genetic perturbations do not affect the core topology of well-established central metabolic enzymes, adaptive evolution, especially when driven by mutations in pleiotropic genes, can impact both the topology of metabolic networks and the identity of the affected metabolites. In our study, mutations in *pntA*, *prfC*, and *rpoB* (**Table 1**) likely produced widespread perturbations in the metabolic network, triggering shifts in global metabolic states, activating latent pathways, and redistributing fluxes across pathways. For example, loss-of-function mutations in *pntA* are known to alter the NADPH/NADP^+^ ratio, influencing numerous cellular reactions^35^. This shift could thermodynamically favor the reverse direction of certain NADP(H)-dependent reactions^80^ and modulate (inhibit or activate) enzymes in key branch points of the metabolic pathways^35, 81^. Such systemic changes would necessitate adjustments to the kinetic and thermodynamic metabolic models used by the MFA and expanding the scope of targeted metabolite detection^79^. This is where metabolite correlation networks offer a key advantage: they allow detecting mutation-induced changes in metabolism without requiring prior knowledge of the exact topological alterations or the identity of affected metabolites. The detected changes offer testable mechanistic hypotheses that can be further validated by complementary tools, such as flux balance analysis and transcriptomics.

We demonstrate the applicability of the proposed framework by mimicking an evolutionary scenario where adaptation of bacterial populations is triggered by an inefficient enzyme. By substituting the *E. coli metK* gene encoding an essential MAT with its ortholog from *U. urealyticum* and allowing the resulting maladapted strain to evolve, we captured snapshots of the evolutionary process through the construction of metabolite correlation networks. We note that correlation networks were constructed only for a single representative variant in the evolved populations. It is plausible, therefore, that the specific mutations and network properties observed in these single isolates might not represent the full diversity of the adaptive changes within the entire evolved populations.

Laboratory evolution disrupted network density, eliminated hub nodes, and weakened correlations among metabolites. However, the consequences of link loss on the ‘small-world’ properties of these networks depended heavily on the structure of the naïve network prior to evolution. When the wild-type (WT) strain underwent laboratory evolution, the global clustering coefficient in the resulting ‘evolved-WT’ network remained largely unchanged. In contrast, both the average shortest path length and the network diameter increased, indicating a reduction in global communication efficiency between network modules without disrupting high node connectivity within modules. Assuming that at least some nodes within modules represent metabolites with shared metabolic functions or regulatory relationships, the preserved local connectivity in the ‘evolved-WT’ network may reflect an essential structural feature that maintains core metabolic functionality. Efficient inter-module communication is likely linked to regulatory mechanisms that coordinate rapid metabolic adjustments to environmental changes. However, because the evolution of the ‘evolved-WT’ strain occurred under constant environmental conditions, the observed reduction in global communication efficiency may have been selectively neutral (or even advantageous) by eliminating unnecessary regulation and potentially improving biomass production efficiency. Thus, although local clustering was maintained, the transformation from the WT to the ‘evolved-WT’ network was accompanied by a loss of ‘small-world’ properties, likely due to the removal of hub nodes that previously served as shortcut connectors across network modules. Additionally, the pronounced (45%) reduction in negative correlations in the ‘evolved-WT’ network, an effect not observed in either the unevolved or the maladapted (’evolved’) networks, may signal a broader loss of metabolic regulation, given the proposed role of negative correlations in regulatory control^25^.

In sharp contrast to the WT network, the ‘unevolved’ network generated by orthologous replacement of the endogenous *metK* gene was markedly disintegrated, exhibiting severely reduced local and global connectivity. Despite a further overall loss of links during the evolution, the resulting ‘evolved’ network regained or even enhanced its ‘small-world’ properties. This observation suggests that evolution does not simply aim to maximize or minimize network connectivity but instead rewires connections in ways that enhance metabolic efficiency within the context of a specific genetic background and selection pressure. Notably, while the evolution of the WT network resulted in the loss of highly connected hub nodes, the evolution of the ‘unevolved’ network led to a more targeted consolidation of connectivity within specific modules, as reflected in an increased global clustering coefficient, and the emergence of strategically placed links that bridged distant modules. These changes point to a distinct evolutionary strategy tailored to the structural constraints of the maladapted starting network. Our findings suggest that the preservation or restoration of ‘small-world’ features that take place despite extensive network rewiring, is a general hallmark of metabolite correlation networks subjected to adaptive selection.

A key example of how metabolite correlation network reorganization relates to fitness is the emergence of S-adenosylmethionine (SAM) as a critical connector node with high betweenness centrality in the ‘evolved’ network. This reorganization was not recapitulated in the ‘mut’ strain, which carries only the *metK* promoter mutation. The increased contribution of the SAM node to inter-module connectivity in the ‘evolved’ network may reflect more efficient SAM utilization. This interpretation is supported by the observation that SAM levels are lower in the ‘evolved’ strain compared to the ‘mut’ strain, despite both exhibiting similar methionine levels—suggesting comparable SAM production (**Fig. 1B**). Therefore, while both strains likely produce equivalent amounts of SAM due to the shared *metK* promoter mutation, additional off-target mutation(s) in the ‘evolved’ strain may have conferred a fitness advantage by enhancing the efficiency of SAM utilization.

By identifying additional nodes with significantly altered properties, we uncovered metabolic pathways likely central to adaptation, including purine, methionine, and polyamine synthesis. The involvement of these pathways offers testable mechanistic hypotheses. For example, a shift in the NADPH/NADP^+^ ratio, potentially triggered by a mutation in *pntA,* may redirect folate-derived one-carbon units away from methionine regeneration and toward purine biosynthesis, thereby alleviating a bottleneck caused by the suboptimal orthologous MAT enzyme (see **Supplementary Note II**).

We propose that the developed experimental and analytical framework of constructing and analyzing metabolite correlation networks holds broader applicability for understanding genotype-phenotype-fitness relationships across diverse evolutionary scenarios and biological systems. Given that metabolic systems are critically linked to organismal fitness and that metabolite correlation networks dynamically reflect the system’s physiological state under genetic and environmental changes, this framework could be extended to study adaptation to various selection pressures beyond inefficient enzymes, such as antibiotic resistance, nutrient limitation, or environmental stresses. Furthermore, the ability to identify key connector metabolites like SAM, whose altered network properties reveal adaptive phenotypes, suggests a valuable tool for uncovering the metabolic underpinnings of adaptation in any organism where comprehensive metabolomics data can be generated. This could have significant implications not only for evolutionary biology but also for applied fields like metabolic engineering, where understanding and manipulating metabolic networks is crucial for optimizing production or creating novel biochemical pathways. The observed preservation or restoration of ‘small-world’ network properties under adaptive selection may represent a general principle of metabolic adaptation, warranting further investigation across different systems to test its universality.

## Materials and Methods

### Culture Medium Preparation

Bacterial strains were cultivated in a minimal M9 medium supplemented with casamino acids. The medium contained M9 salts (15 g/L KH_2_PO_4_, 2.5 g/L NaCl, 33.9 g/L Na_2_HPO_4_, 5 g/L NH_4_Cl), casamino acids (0.1%), thiamine (0.5 μg/L), magnesium sulfate (1 mM), and D-glucose (2 g/L). The final volume was adjusted with sterilized water. M9 salts were obtained from Formedium (Swaffham, UK). Thiamine (99%) and Bacto Casamino Acids were purchased from Thermo Fisher Scientific (St. Louis, MO), while D-glucose (≥99.5%) and magnesium sulfate (≥99.5%) were sourced from Sigma-Aldrich (St. Louis, MO). The medium was sterilized by autoclaving.

### Genome editing and Experimental Evolution

In this study, wild-type (WT) *E. coli* K-12-MG1655 strain was employed as the model organism. A replacement of the essential *metK* gene encoding methionine adenosyltransferase (MAT) with an ortholog from *Ureaplasma urealyticum* was conducted in two steps. First, a Tn7 transposon-based system was used to introduce the coding sequence of orthologous *metK* under the control of the endogenous *E. coli metK* promoter into the genome of *E.coli* ^82^. Once the orthologous *metK* gene was successfully integrated, a λ Red recombination system^83^ was used to delete the endogenous *E. coli metK*. The whole genome sequencing validated that the resulting ‘unevolved’ strain was genetically identical to the WT except for the altered gene. Subsequently, the WT and unevolved bacterial populations were subjected to adaptive laboratory evolution by serial passaging. The populations were grown in 96-well microtiter plates and propagated by 1/100 dilution into fresh media every 24 hours. A total of 48 passages produced ‘evolved-WT’ and ‘evolved’ populations. An additional ‘mut’ strain was produced by incorporating a single (-8) T to A *metK* promoter mutation using a λ Red recombination system on the ‘unevolved’ strain genetic background. Each of the listed bacteria strains was stored in a glycerol stock at -80°C for further analysis.

### Growth Rate Evaluation

To measure the growth rates of bacterial strains, a 5 mL starter culture of each strain was grown overnight at 37°C in the culture medium. The overnight cultures were subsequently diluted at 1:100, and 50 mL of each diluted culture was incubated under identical conditions until reaching an optical density (OD) at 600 nm of 0.5. The cultures were then further diluted 1:50 to the same final volume and incubated for an additional 10 hours at 37°C. Optical density was measured and recorded every 30 minutes using an Infinite 200 PRO plate reader (Tecan, Switzerland) at 600nm. To compare the growth rates of the bacterial strains, we fitted the growth data using a logistic model^84^:

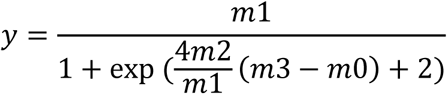

In this model, *m*1 represents the asymptotic maximum (A), *m*2 corresponds to the maximum specific growth rate (μ max), and *m*3 indicates the lag phase duration (λ). To statistically compare the different parameters between strains, we performed a one way ANOVA, followed by pairwise comparisons using Tukey’s HSD.

### Quantitative Western Blot Analysis of MAT Levels

A quantitative Western Blot was performed to assess the relative expression levels of methionine adenosyltransferase (MAT). Cells were lysed using a lysis buffer (25 mM Tris, pH 8.0; 150 mM KCl; 1 mM BTT; 1% Triton X-100 (Sigma); 0.5% Benzonase (Sigma)) after a 15-minute pre-incubation. The samples were diluted in Tris buffer (pH 8.0) to ensure that 10 μg of protein was loaded per well onto a 12.5% SDS-PAGE gel. Electrophoresis was conducted at 120 V for 80 minutes, followed by electrotransfer onto a Trans-Blot Pure nitrocellulose membrane (Bio-Rad, Hercules, CA). The membrane was blocked with AdvanBlock-PF blocking solution (Advansta, San Jose, MO) and washed with AdvanWash washing solution (Advansta). Custom-raised rabbit anti-*Ureaplasma urealyticum* MAT polyclonal antibodies (Genemed Synthesis, San Antonio, TX) were added, and the membrane was incubated for one hour at room temperature with shaking. After additional washes with AdvanWash, the membrane was incubated with goat anti-rabbit horseradish peroxidase (HRP)-conjugated secondary antibodies (SH026; Abm) for one hour at room temperature. The immunoblot signal was detected using enhanced chemiluminescence (ECL; Advansta). The relative intracellular abundances of methionine MAT were compared using pairwise t-tests with Bonferroni correction.

### Whole Genome Sequencing

A comprehensive whole genome sequencing (WGS) analysis was conducted to obtain genomic data for the ‘unevolved’, ‘evolved’, ‘evolved-WT’, and ‘mut’ strains, which were then compared to the genome sequence of the WT strain. DNA extraction was performed using the DNeasy UltraClean Microbial Kit (QIAGEN, Hilden, Germany) according to the manufacturer’s protocol. Sequencing was carried out by the Mantoux Bioinformatics team at the Ilana and Pascal Mantoux Institute for Bioinformatics, Weizmann Institute, Israel. The MiSeq Micro V2 (300 cycles) kit (Illumina, San Diego, CA) was used for sequencing, and library preparation included the incorporation of nine-base Unique Molecular Identifiers (UMIs).

### LC-MS Analysis

Two experimental batches, each consisting of 19 biological replicates per bacterial strain, were prepared, resulting in a total of 38 replicates per strain. Cultures were grown overnight at 37°C in the culture medium, followed by a 1:100 dilution and further incubation until reaching an OD of 0.5 at 600 nm. A secondary 1:50 dilution was then performed, with additional growth until the OD reached 0.5 again. Cells were then harvested and centrifuged at 4 °C for 15min at 4000 × g. The supernatant was discarded, and the pellet was resuspended in 1 ml of ice-cold phosphate-buffered saline and centrifuged again in the same conditions. Afterwards, ice-cold 80% ultrapure methanol was added to the pellet, mixed well by vertexing, put for 10 s in liquid nitrogen, and incubated for 30min at −20 °C. To remove cell debris, samples were centrifuged at a maximum speed at 4 °C for 30min. The supernatant was collected into a fresh Eppendorf tube and by using a SpeedVac vacuum concentrator (Thermo Fischer Scientific) the liquid was evaporated. Pure standards of S-adenosylmethionine (SAM) and methionine were prepared to facilitate their identification in the chromatograms. Metabolite analysis was performed using Acquity UPLC I-Class System (Waters Corporation, Milford, MA) coupled with a Q Exactive hybrid quadrupole-orbitrap mass spectrometer (Thermo Fisher Scientific) in positive ionization mode.

### Screening for Unknown Features using Compound Discoverer

Raw data files were processed using Compound Discoverer version 3.3 SP2 (Thermo Fisher Scientific) to screen for unknown compounds. The analysis workflow included retention time alignment, unknown peak detection and ion association, peak filtration (intensity ≤ 1e6), gap filling, and removal of background components unrelated to experimental samples (detected in blanks with a ratio ≤ 2.5). Additional steps involved elemental composition prediction, database searching via ChemSpider, spectral matching with mzCloud, and mass list comparisons. To avoid redundant identification of multiple adduct ion compositions of the same compound, metabolic features were consolidated based on similarity in retention time (RT difference ≤ 0.1), high correlation (R ≥ 0.9), and m/z differences matching known adducts. The program annotations of SAM and methionine were validated using pure standards, and their levels across strains were compared using one-way ANOVA, followed by pairwise comparisons using Tukey’s HSD test.

### Principal Components Analysis (PCA) of the metabolites

A two-dimensional principal components analysis (PCA) was performed using MetaboAnalyst 5.0, a freely accessible web-based platform for metabolomics data analysis, statistical evaluation, and data visualization (MetaboAnalyst). Prior to analysis, the data were log-normalized.

### Construction of Correlation Networks

Correlation analysis was used to evaluate associations between variables in the untargeted LC-MS metabolomics results. Pearson’s pairwise correlations were computed based on the peak intensities of metabolic features, generating an adjacency matrix of correlation coefficients. To remove transitive interactions, we employed the Probabilistic Context Likelihood of Relatedness (PCLRC) algorithm, an extension of the Context Likelihood of Relatedness (CLR) approach^63, 85^. PCLRC integrates correlation analysis with mutual information and resampling techniques to filter out non-significant associations. In each iteration, 75% of the data is randomly sampled, correlations are computed, and CLR is applied. Only the top Q% of interactions are retained, and after k iterations, edges present in at least 95% of iterations form the final network. Following established recommendations^24, 63^, we set Q% to 0.3 and k to 1,000.

For network construction, metabolic features were grouped by bacterial strain and experimental batch. An undirected, weighted network, in which edge weights represented the magnitude of correlations, was generated for each group, retaining only consistent correlations across technical replicates. This approach improved network reliability by minimizing noise and random variation.

### Network analysis

*Average Shortest Path* and *Network Diameter* were calculated with ‘mean_distance’ and ‘diameter’ functions, respectively, using ‘igraph’ package in R^86^. Weights parameter was set to 1/(correlation coefficient of the edges) to account for edge strength, treating higher weights as closer connections in the network.

*Node degree* was calculated with ‘degree’ function from the ‘igraph’ package in R^87^. *Modularity* was calculated by applying ‘Infomap’ algorithm from ‘infomapecology’ package in R^87^ on a weighted bilayer network composed of layers of positive and negative correlations. In this approach, each metabolic feature (“physical node”) is represented by two “state nodes” in the network—one in the positive layer and one in the negative layer—enabling the analysis of both types of correlations within the same framework. These two state nodes may not belong to the same module, and they may not share similar characteristics.

*Normalized Mutual Information (NMI)* was calculated using ‘NMI’ function in ‘infomapecology’ package in R^87^. This measure allows the comparison of two partitions even when they have a different number of modules. Given two network partitions, the NMI is calculated as^88^:

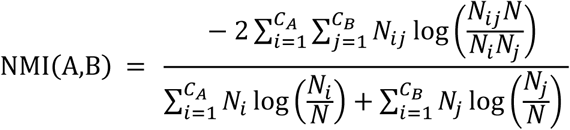

Where *A* and *B* are the two network partitions; *C_A_* is the number of modules in partition *A*, *C_B_* is the number of communities in partition *B*, *N* is the number of nodes, N_ij_ is the overlap between *A*’s module *i* and *B*’s module *j*, *i.e.*, the number of nodes that the two modules have in common. *N_i_* is the total number of nodes in *A*’s module *i*, and N*j* is the total number of nodes in *B*’s module *j*. The NMI ranges from 0 to 1, where 0 signifies that the community structures are independent and 1 that they are identical.

*Neighborhood Connectivity* was calculated using the ‘neighborhood.connectivity’ function from the ‘influential’ package in R^89^:

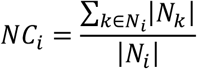

Where *N_i_* is the set of neighbors of vertex *i*, and *k* is one of these neighbors.

*Clustering Coefficient* was calculated using ‘transitivity’ function from the ‘igraph’ package in R^86^. It is defined as the number of triangles connected to a node *i* divided by the number of triples centered around it:

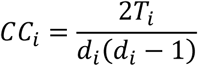

where *T_i_* is the count of triangles connected to node *i*, and *di* is the degree value of node *i*. *Global Clustering Coefficient* was calculated as the ratio of the count of triangles and connected triples in the graph.

*Node Role* was determined by its within-module connectivity and its participation across different modules, both of which reflect the node’s function and position in the network^72^. The classification of node roles was based on two key parameters—the within-module degree (*Z_i_*) and the participation coefficient (*C_i_*). *Z_i_* measures how well-connected a node is within its own module. Nodes with high *Z_i_* values are considered “module hubs” because they have strong connections within their module. In contrast, nodes with low *Z_i_* values have weaker connections within their module and are categorized as “non-hubs.” *C_i_* evaluates how a node’s connections are distributed across different modules. A *C_i_* value close to 1 suggests that the node connects to many modules, while a value close to 0 implies the node is confined to its own module. This is used to further distinguish between roles: Peripheral nodes are non-hubs with a low *C_i_* (*C_i_* ≤ 0.62), indicating that their connections are mainly within their own module; Connectors are non-hubs with a high *C_i_* (*C_i_* > 0.62), meaning their connections span multiple modules; Module hubs are hubs with a low *C_i_* (*C_i_* ≤ 0.62), as they are highly connected within their module but don’t bridge different modules; Network hubs are hubs with a high *C_i_* (*C_i_* > 0.62), indicating that they connect different modules and play a central role in linking various parts of the network.

### Targeted Metabolomics

To advance the identification of unknown compounds based on fragmentation spectral information, we used SIRIUS 6.1.0 software^73^. The isotope patterns of the molecular ions from the MS1 and the fragmentation spectra from the MS2 of the selected compounds were manually exported from CompoundDiscoverer and imported into SIRIUS. The following parameters were used for the computation: instrument (Orbitrap), MS2 mass accuracy (5 ppm), possible adducts ([M+H]+, [M+Na]+, [M+K]+), molecular formula generation (de novo + bottom up). In addition, CSI:FingerID fingerprint prediction^75^, CANOPUS compound class identification^74^, CSI:FingerID structure database search, and MSNovelist^76^ were activated, all other parameters were maintained at their default settings.

## Supplementary Information

**Figure S1.**
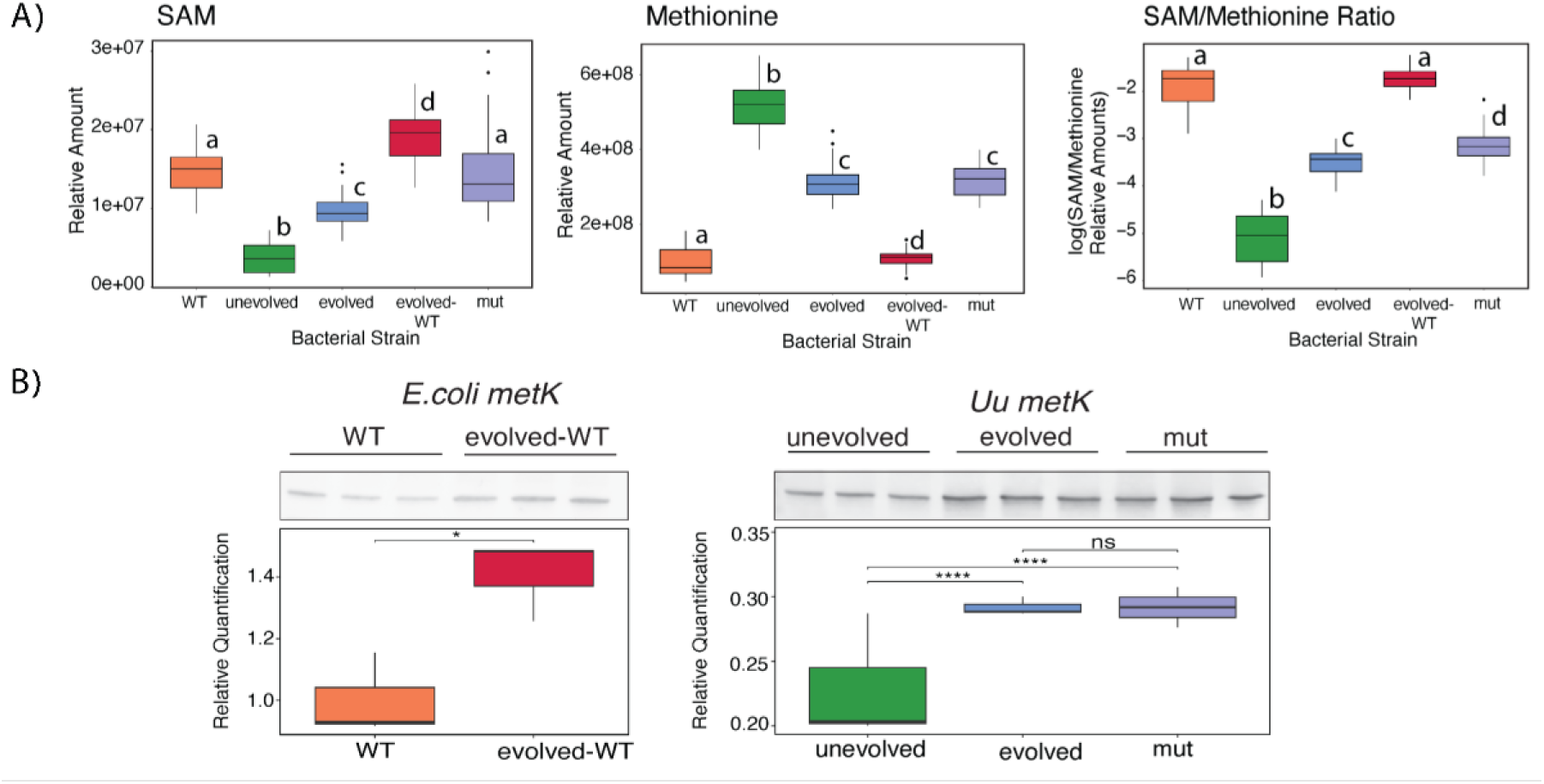
**A.** Relative intracellular abundance and abundance ratios. *Left panel*, S-adenoslmethionine (SAM); *middle panel*, methionine in the engineered and evolved strains as detected by targeted LC-MS/MS analysis. *Right panel*: ratios between SAM and methionine levels. Following one-way ANOVA, Tukey HSD post-hoc analysis assigned letters (a, b, c, d) to indicate significant strain differences (*p-values*<0.05). Strains not sharing a letter exhibit a statistically significant mean difference in the relative abundance. **B**. Western Blot analysis of relative intracellular abundance of methionine S-adenosyltransferase (MAT) following gene editing and evolution. * *p-value* <0.05; *** *p-value* <0.001, ns – not significant (Pairwise *t*-test with Bonferroni adjustment).

**Figure S2.**
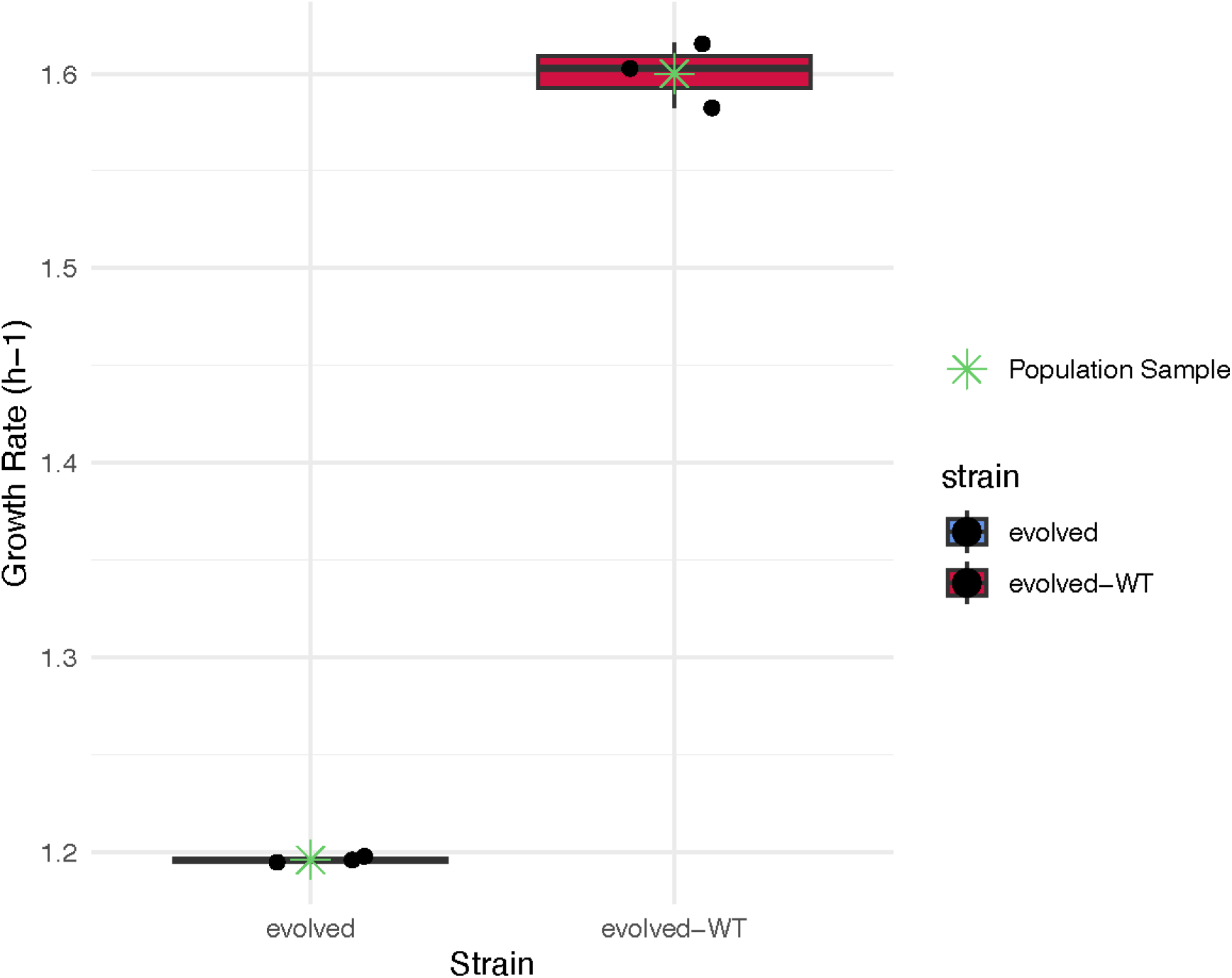
Comparisons of growth rates of variants isolated from the ‘evolved’ and ‘evolved-WT’ populations. Black dots correspond to randomly isolated variants *v1-v3*.

**Figure S3.**
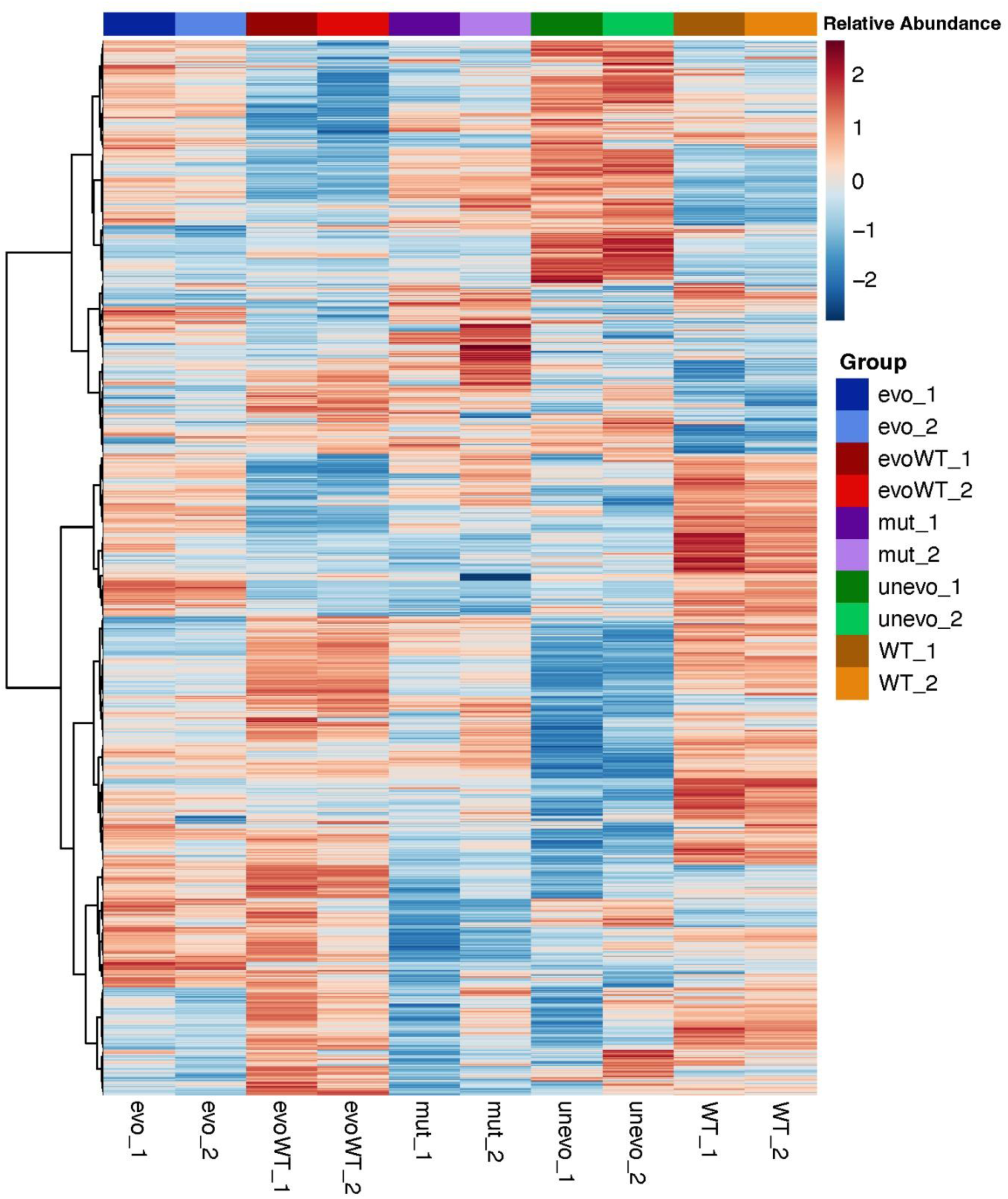
Heat map of relative untargeted metabolite abundances. Both batches for each strain are shown.

**Figure S4.**
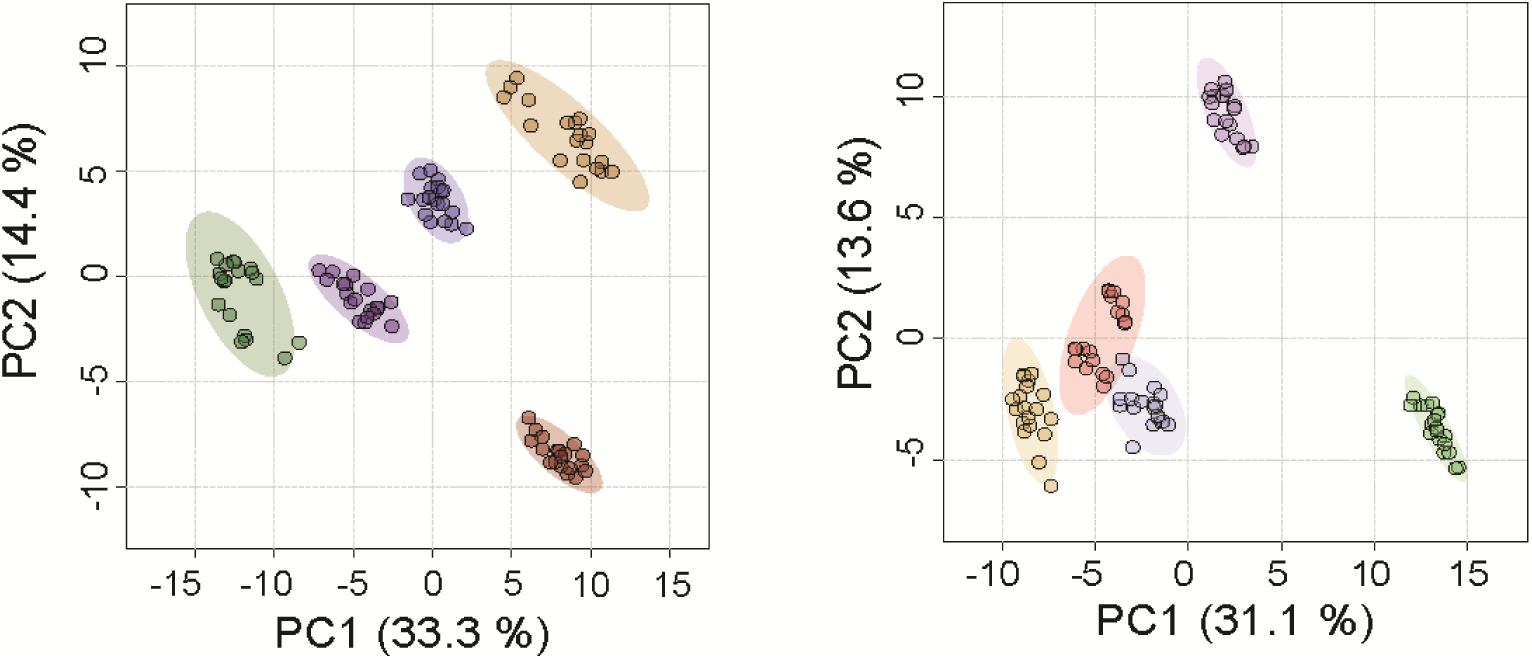
Principal component analysis (PCA) of the relative abundances of untargeted metabolic features extracted from both batches. Each data point represents a biological replicate within a batch. The *left* and *right* panels represent batches 1 and 2, respectively.

**Figure S5.**
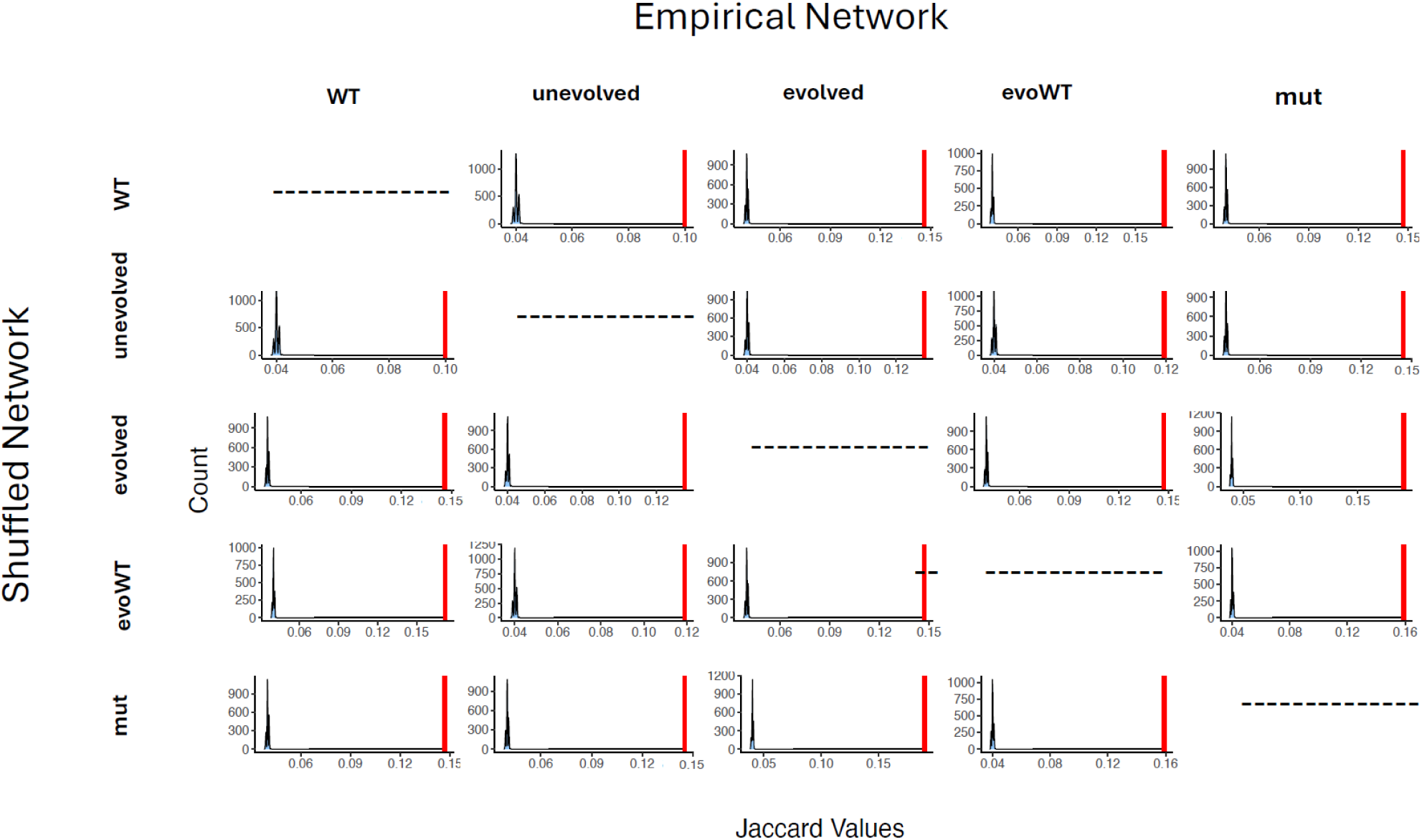
Jaccard similarity values of empirical and shuffled networks. The red vertical lines represent the empirical Jaccard similarity values as summarized in Fig. 1E, and the blue curves represent the distribution of the values of the comparisons to the random networks. All Jaccard similarity values display clear statistical significance (*p-value* << 10^-4^, one-sample *z*-test).

**Figure S6.**
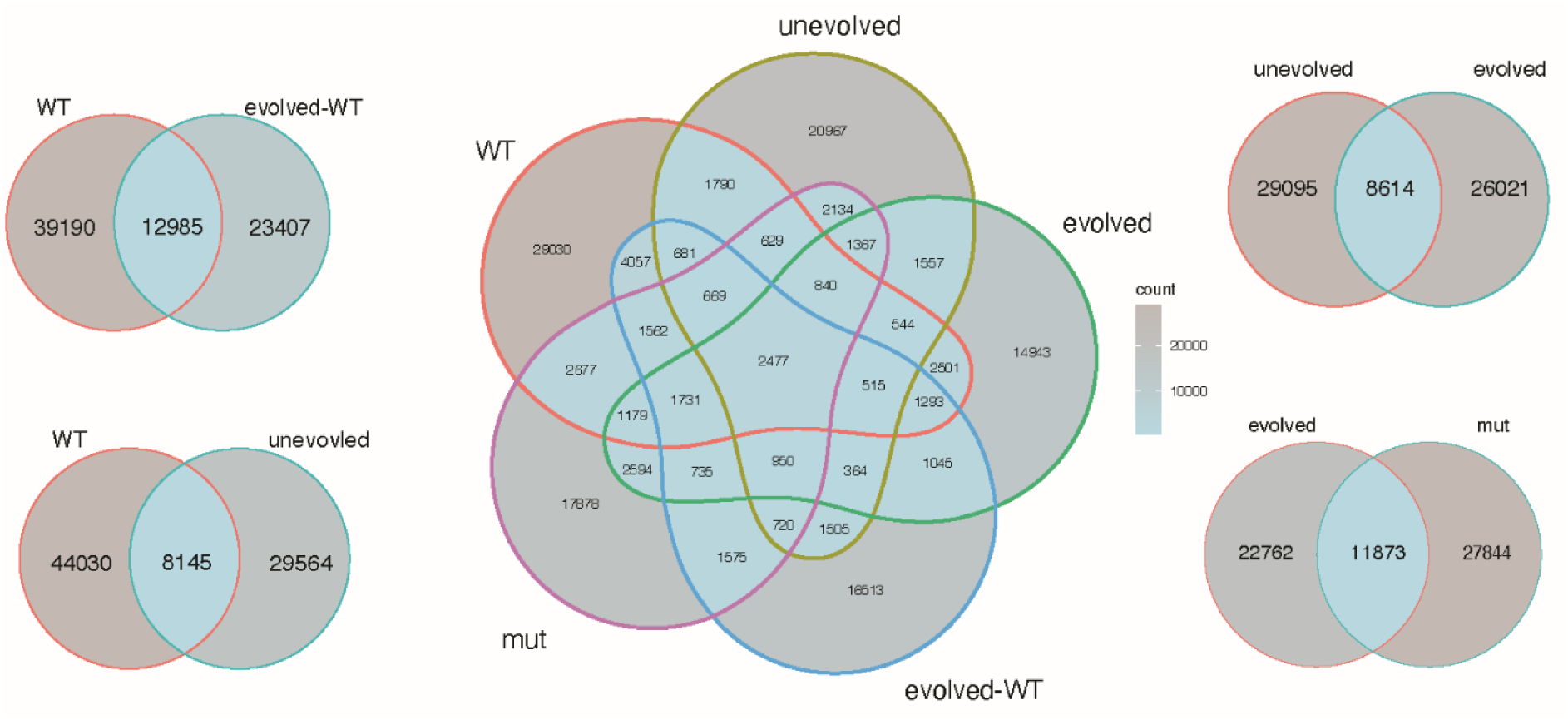
Shared correlations (both positive and negative) between strains.

**Figure S7.**
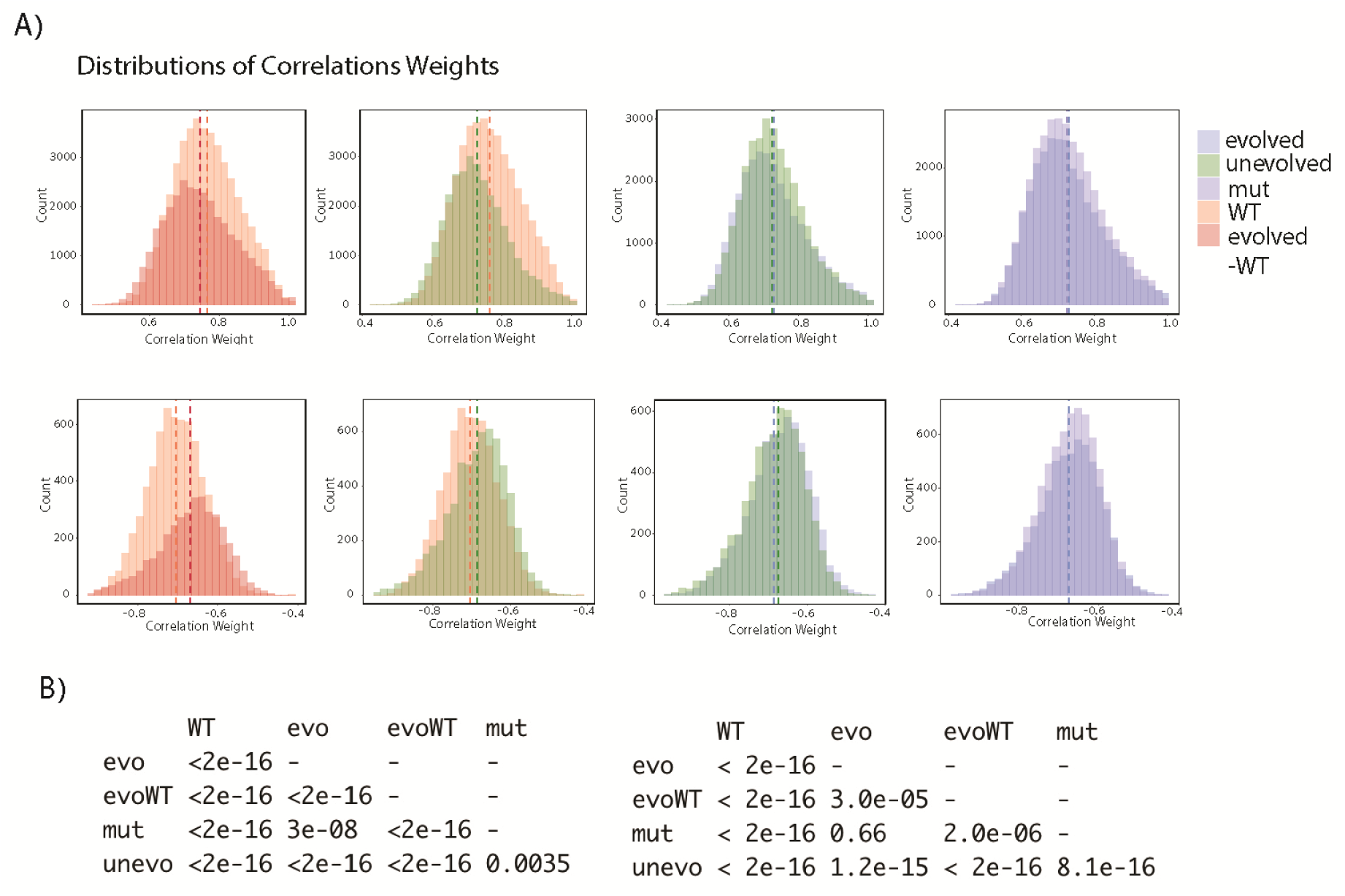
**A.** Distribution of correlation weights. *Upper panel*, positive correlations; *lower panel*, negative correlations. **B**. *p-values* of pairwise comparisons using Wilcoxon rank trust sum test with BH-FDR corrections. *Left panel*, distributions of positive correlations. *Right panel*, distributions of negative correlations.

**Figure S8.**
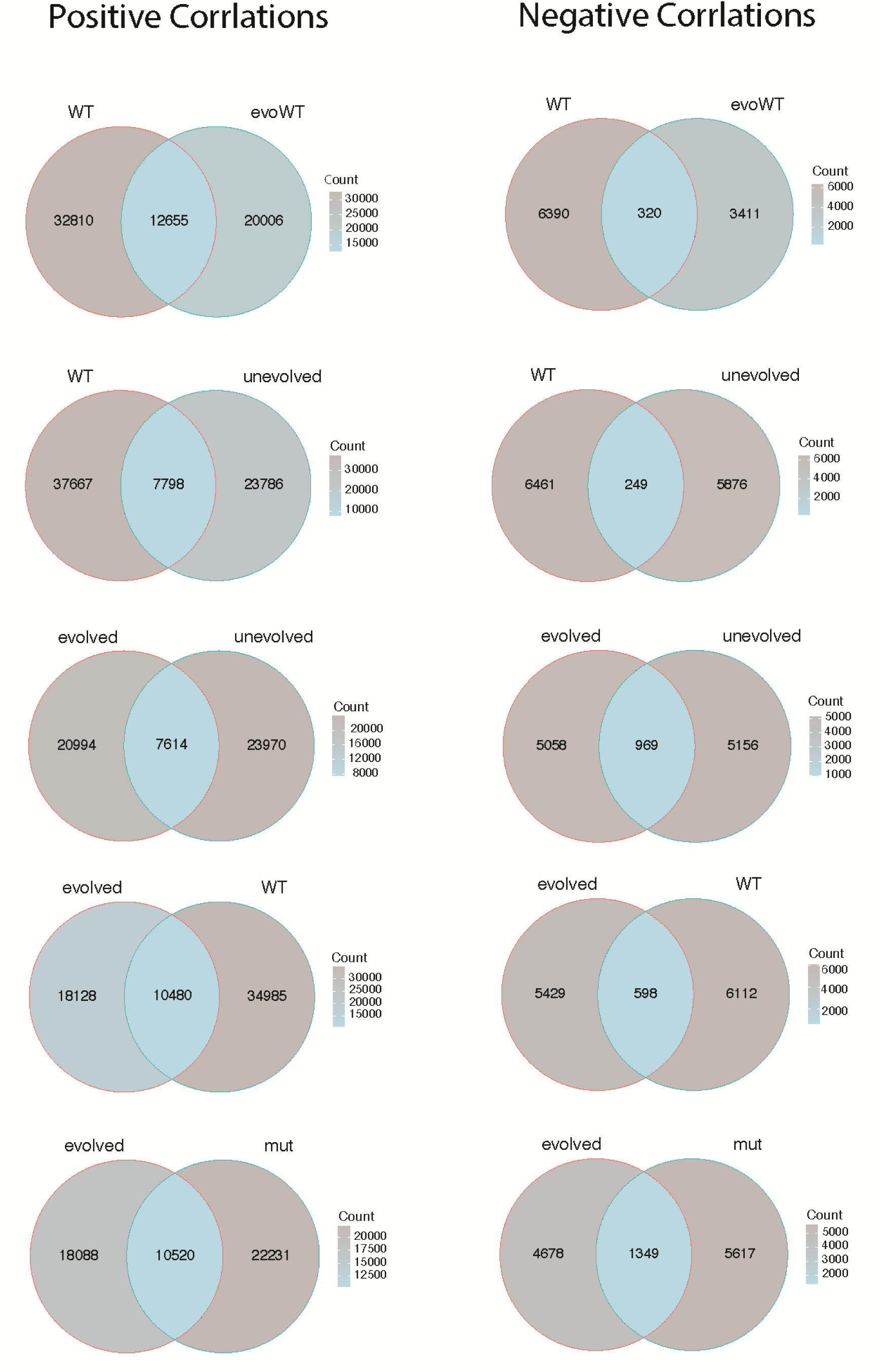
Positive and negative shared correlations between strains.

**Figure S9.**
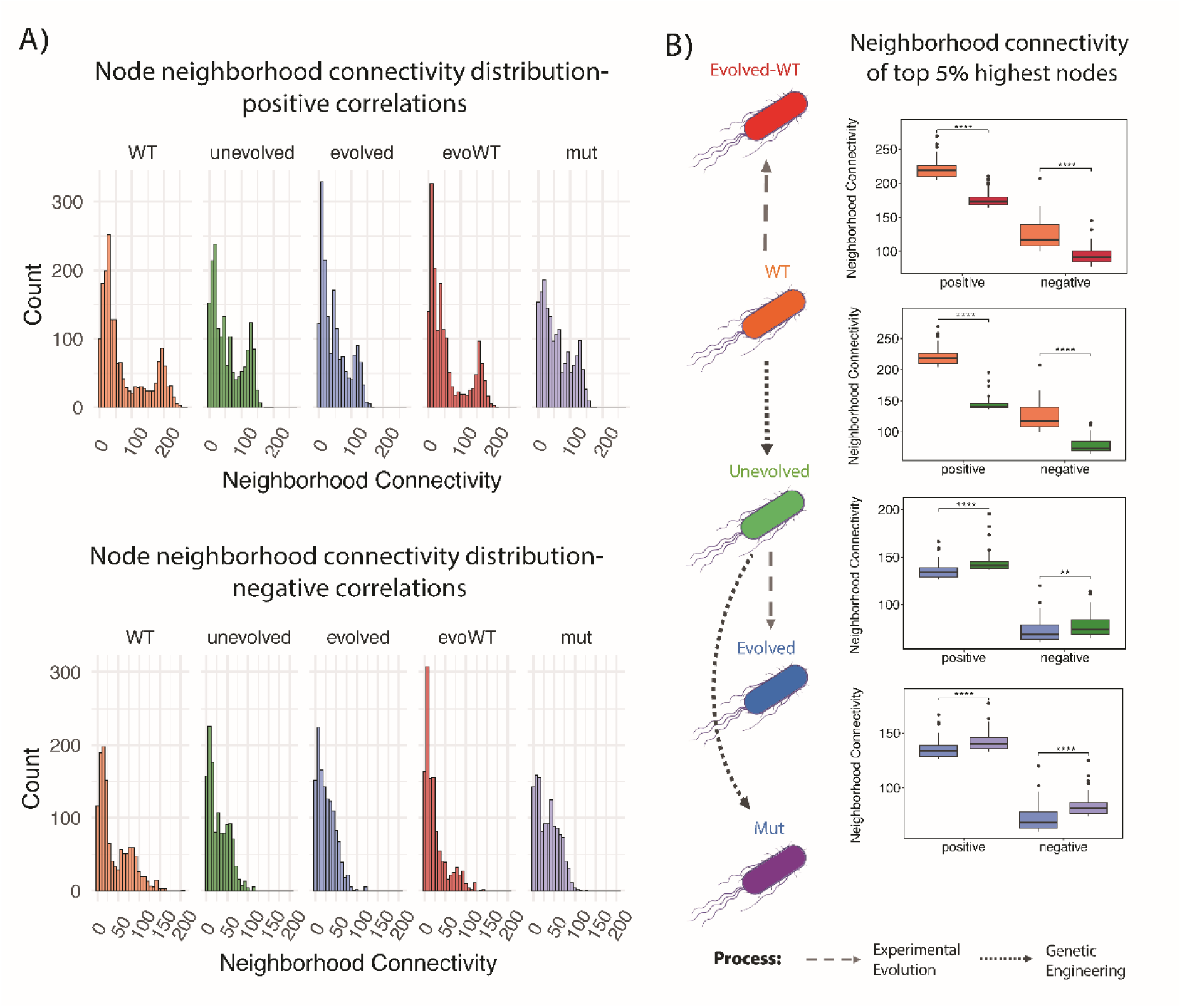
**A**. Network Neighborhood Connectivity (NC) distributions. *Upper panel* – positive correlations. *Lower panel* – negative correlations. **B**. *Left panel*, color coding of network strains. *Right panel*, comparisons of neighborhood connectivity (NC) between top 5% nodes with highest NC in each network. **** *p-value* <10^-4^, *** *p-value* <10^-3^ (two-sample *t*-test); ns – nonsignificant.

**Figure S10.**
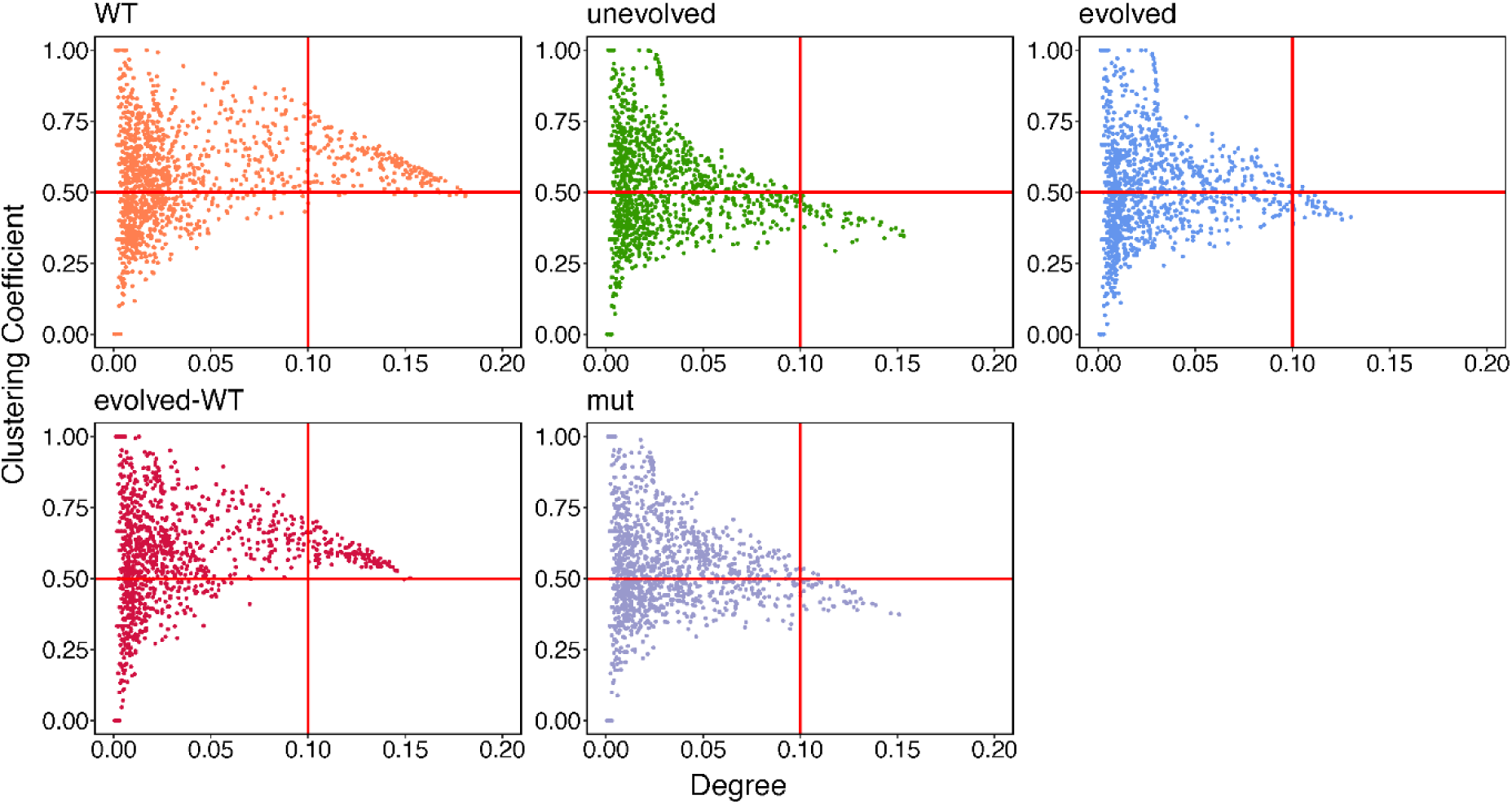
Normalized node degree as a function of clustering coefficient.

## Supplementary Note I

In a regular network, GCC is independent of network size and equals ∼ 0.75; *L* is around *n*/2*k*, where *n* is the number of nodes and *k* is the average degree^1^. For a regular network, in which all nodes have the same degree, with *n* and *k* values identical to those of the WT network (that is, *n* = 2,118 nodes and each node has a degree of *k* = 52,175 links/2,118 nodes = 24.6), *L* is approximately *n/2k ∼* 43. Conversely, in a random network (in which the probability of a link is uniform), GCC can be estimated as *k*/*n*, and *L* typically scales with ln(*n*)/ln(*k*)^1^. For a random network of WT network size with *n =* 2,118 nodes and average degree *k* = 24.6, GCC would be around 0.012 and *L* around 2.39. Based on the finding that the GCC value of the WT network is substantially higher than what is typically seen in a random network (0.59 vs. 0.012, **Table 2**), and that the average shortest path *L* is much smaller than expected in a regular network (5.74 for positive correlation vs. the expected 43, **Table 2**), we conclude that the metabolite correlation network of the WT strain exhibits “small-world” properties.

## Supplementary Note II

Out of 18 identified metabolites, five participate in purine biosynthesis (**Table S5**). Specifically, L-glutamine and SAICAR serve as a substrate and an intermediate, respectively, in the purine biosynthetic pathway^1^. Xanthosine functions as a precursor in the purine salvage pathway^2, 3^. 5,10-Methenyltetrahydrofolate is a direct precursor of 10-formyltetrahydrofolate, which donates one-carbon units required for purine synthesis^4^. S-hydroxymethyl glutathione acts as a precursor of formate^5^, which contributes to an alternative route for the production of 10-formyltetrahydrofolate^6^.

What might explain the changes in the network properties of nodes representing these metabolites? SAM biosynthesis competes with the purine biosynthetic pathway for one-carbon units supplied by the folate cycle. A key branching point in this cycle directs 5,10-methylenetetrahydrofolate either toward 5-methyltetrahydrofolate, which is used to regenerate methionine for SAM synthesis, or toward 5,10-methenyltetrahydrofolate, a precursor of 10- formyltetrahydrofolate that donates one-carbon units for purine biosynthesis. This branch point is regulated by the intracellular NADPH to NADP^+^ ratio^7, 8^. The one-carbon unit transferred toward methionine synthesis is recycled primarily through the activity of MAT. When MAT function is compromised and SAM levels are low, one-carbon units may become sequestered in methionine, without an efficient route for reutilization in other biosynthetic processes. Under these conditions, redirecting folate flux toward 10-formyltetrahydrofolate and enhancing purine biosynthesis may offer a selective advantage. This metabolic reorganization could be driven by the loss-of-function mutation in the *pntA* gene observed in the evolved strain (**Table 1**). Furthermore, limiting the methionine biosynthesis pathway could prevent methionine accumulation and help alleviate the associated metabolic bottleneck. Consistent with this model, another identified metabolite, L-threonine may play a regulatory role through its known ability to inhibit de novo methionine biosynthesis^9^. (We also identified L-homoserine phosphate that is a metabolic precursor of L-threonine).

Another three identified metabolites—palmitoylputrescine, N acetylputrescine, and methylthioadenosine—are associated with polyamine biosynthesis. Polyamines, derived from the aminopropyl group of SAM, play essential roles in bacterial growth and physiology^10^. Shifts in the network properties of nodes representing these metabolites in the ‘evolved’ strain may reflect a more efficient use of SAM, with metabolic fluxes utilizing SAM redirected toward pathways that are critical for cellular growth.

Finally, three additional metabolites—enterobactin, cyclic AMP (cAMP), and γ- glutamylcysteine—play global roles in *E. coli* metabolism and may be responsible for the profound changes observed in the properties of the ‘evolved’ network. Enterobactin is a siderophore that promotes growth by facilitating iron uptake^11^. cAMP acts as a secondary messenger that activates the cAMP receptor protein (CRP), a transcription factor that regulates the expression of more than 180 catabolic genes. The cAMP-CRP signaling pathway supports bacterial growth by coordinating the expression of catabolic genes with the availability of metabolites^12^. Gamma glutamylcysteine is the immediate precursor of glutathione, an essential molecule for protection against oxidative stress^13^.

